# Amperometric measurements of cocaine cue and novel context-evoked glutamate and nitric oxide release in the nucleus accumbens core

**DOI:** 10.1101/826677

**Authors:** BM Siemsen, JA McFaddin, K Haigh, AG Brock, MN Leath, KN Hooker, LK McGonegal, MD Scofield

**Affiliations:** Department of Anesthesia and Perioperative Medicine, Medical University of South Carolina, Charleston, SC, USA; Department of Neuroscience, Medical University of South Carolina, Charleston, SC, USA

**Keywords:** glutamate, nitric oxide, cocaine, reinstatement, nNOS, novelty

## Abstract

Cue-induced reinstatement of cocaine seeking after self-administration (SA) and extinction relies on glutamate release in the nucleus accumbens core (NAcore), which in turn activates neuronal nitric oxide synthase (nNOS) interneurons. Nitric oxide (NO) is required for structural plasticity in NAcore medium spiny neurons (MSNs), as well as cued cocaine seeking. However, NO release in the NAcore during reinstatement has yet to be directly measured. Further, the temporal relationship between glutamate release, and the induction of a NO response also remains unknown. Using wireless amperometric recordings in awake behaving rats, we quantified the magnitude and temporal dynamics of novel context- and cue-induced reinstatement-evoked glutamate and NO release in the NAcore. We found that re-exposure to cocaine-conditioned stimuli following SA and extinction increased extracellular glutamate, leading to release of NO in the NAcore. In contrast, exposing drug-naïve rats to a novel context led to a lower magnitude rise in glutamate in the NAcore relative to cue-induced reinstatement. Interestingly, novel context exposure evoked a higher magnitude NO response relative to cue-induced reinstatement. Despite differences in magnitude, novel context evoked-NO release in the NAcore was also temporally delayed when compared to glutamate. These results demonstrate a dissociation between the magnitude of cocaine cue- and novel context-evoked glutamate and NO release in the NAcore, yet similarity in the temporal dynamics of their release. Together, these data contribute to a greater understanding of the relationship between glutamate and NO, two neurotransmitters implicated in encoding the valence of distinct contextual stimuli.

## 1. Introduction

Persistent relapse vulnerability represents a significant hurdle in the clinical treatment of addiction (Sinha 2011). To better inform the development of novel therapeutics, cue-induced relapse has been extensively modeled in the laboratory as means to disentangle the neurobiological underpinnings of drug craving and relapse (Kalivas 2008; Epstein *et al*. 2006). Preclinical rodent models of cocaine self-administration (SA) and cue-induced reinstatement (Bossert *et al*. 2013) demonstrate that susceptibility to relapse is mechanistically linked to dysfunction in glutamatergic neurotransmission within the nucleus accumbens (Kalivas & Volkow 2011; Scofield *et al*. 2016a). Indeed, *in vivo* microdialysis data demonstrate that both relapse precipitated by conditioned cue exposure, and by a priming injection of cocaine, evoke a pronounced increase in glutamate release in the nucleus accumbens core (NAcore), deemed ‘glutamate overflow’ (Scofield *et al*. 2016a). Mechanistically, glutamate overflow during cocaine seeking occurs due to decreases in basal extrasynaptic glutamate levels following exposure to cocaine (Knackstedt *et al*. 2010). Cocaine-induced decreases in extrasynaptic glutamate levels limit the activation of presynaptic release-regulating mGluR2/3 autoreceptors (Knackstedt *et al*. 2010), which potentiates synaptic glutamate release during cue exposure (Scofield *et al*. 2016a). In addition, cocaine-induced downregulation of the astrocytic glutamate transporter (GLT-1)(Reissner *et al*. 2015), and retraction of astroglial processes from synapses in the NAcore (Scofield *et al*. 2016b) also act in concert to facilitate cue-induced glutamate spillover and drug seeking.

Downstream of cue-induced glutamate release, dysfunctional synaptic plasticity at excitatory synapses in the NAcore is also required for relapse (Moussawi *et al*. 2011). Specifically, cue-induced glutamate release engages a transient synaptic potentiation (t-SP) in NAcore medium spiny neurons (MSNs); increasing AMPA/NMDA ratios and dendritic spine head diameter (d_h_) in a manner that positively correlates with the magnitude of drug seeking (Gipson *et al*. 2013). The structural plasticity component of t-SP requires cue-induced activation of matrix metallo-proteinase (MMP) and degradation of the extracellular matrix (ECM) proteins, allowing for the physical remodeling of dendritic spines and insertion of AMPA receptors (Smith *et al*. 2014). We have recently demonstrated that MMP activation during cue-induced reinstatement occurs via S-nitrosylation of latent pro forms of MMP molecules (Smith *et al*. 2017). S-nitrosylation is a post-translational modification engaged by the gaseous neurotransmitter nitric oxide (NO) (Gu *et al*. 2002). NO can be generated from several sources (Rojo *et al*. 2014; Forstermann & Sessa 2012), yet is primarily produced in the striatum by a sparse population of interneurons that express the Ca^2+^-sensitive enzyme neuronal nitric oxide synthase (nNOS) (Centonze *et al*. 1999). Using anesthetized amperometric NO recordings, we have previously demonstrated that activation of NAcore nitrergic interneurons using Gq-coupled Designer Receptors Exclusively Activated by Designer Drugs (DREADDs) evokes NO release *in vivo*. In addition, selective chemogenetic activation of nNOS interneurons in the NAcore potentiates or induces cocaine seeking in the presence or absence of conditioned cues, respectively (Smith *et al*. 2017). Further, puff application of the mGluR5 agonist CHPG into the NAcore during anesthetized recordings also evokes NO release. Finally, microinfusion of CHPG into the NAcore was sufficient to induce cocaine seeking, an effect that was prevented by prior pharmacological inhibition of nNOS (Smith *et al*. 2017). Taken together, these data suggest that cue-induced glutamate release leads to activation of glutamate receptors on nNOS interneurons, and that glutamate-mediated activation of nNOS neurons is required for NO release and relapse. While evidence clearly suggests a link between cue-induced glutamate release, activation of nNOS interneurons, NO release, and relapse, quantitative assessment of NO release in awake animals has not been conducted. Further, the temporal relationship between cue-induced glutamate and NO release in the NAcore remains to be elucidated.

*In vivo* microdialysis and HPLC-based analytical techniques have been especially useful in delineating the involvement of elevated extracellular NAcore glutamate in cocaine seeking (McFarland *et al*. 2003; Kau *et al*. 2008; Park *et al*. 2002). However, electrochemical-based approaches have a clear advantage over microdialysis with respect to temporal resolution (Burmeister & Gerhardt 2001). Further, microdialysis cannot be used to directly measure NO, with only metabolites of NO production able to be quantified with this technique (Goren *et al*. 2001). Recently, methods for electrochemical-based detection of glutamate and NO have been adapted for wireless data collection, allowing for electrochemical recordings in freely-moving, behaving rats (Hascup *et al*. 2010; Rutherford *et al*. 2007). With this technique it has been demonstrated that cue-induced reinstatement of ethanol-, but not food-, seeking elevates extracellular glutamate in both the basolateral amygdala (BLA) and NAcore (Gass *et al*. 2011). Our lab and others have used specially designed electrochemical sensors to detect NO in brain tissue (Friedemann *et al*. 1996; Smith *et al*. 2017), yet to date only in anesthetized animals (Santos *et al*. 2011; Barbosa *et al*. 2008). These studies demonstrate a relationship between glutamate receptor activation and NO release, with microinfusion of glutamate or NMDA receptor agonists into the CA1 region of the hippocampus transiently elevating NO in anesthetized rats (Barbosa *et al*. 2008). However, to our knowledge, the application of NO recordings in awake and behaving rats has yet to be described. Here we quantify and analyze the temporal relationship between glutamate and NO release in the NAcore during cue-induced reinstatement, comparing these results with those obtained during exposure to a novel environment with interactable levers.

## Materials

### Drugs

DETA-NONOate (DETA) was purchased from Cayman chemicals (Ann Arbor, MI) and was dissolved in NaOH at a concentration of 10 mM for *in vitro* calibrations as well as during *in vivo* recordings. 7-Nitroindazole (7-NI) was purchased from TCI America (Portland, OR). 7-NI was dissolved in DMSO at a concentration of 400 mM. Clozapine-N-oxide (CNO) was obtained through the National Institute of Mental Health Chemical Synthesis Program. CNO was dissolved in 10% DMSO in sterile saline and was delivered at 5 mg/kg (i.p.).

### Biosensors

Glutamate oxidase (GluOx)-coated biosensors (Gass *et al*. 2011; Hu *et al*. 1994) were purchased from Pinnacle Technology (Lawrence, KS). Biosensors consisted of a PTFE-coated electrode (240 µm O.D.), housing a Pt/Ir sensing cavity (176 O.D., 1 mm length) with an epoxy tip at the end of the electrode (200 µm O.D.). Electrodes contained an integrated Ag/AgCl reference electrode (350 µm). Each glutamate electrode was purchased pre-coated with an enzyme layer containing GluOx and ascorbate oxidase (AscOx). GluOx catalyzes the breakdown of glutamate into α-ketoglutarate and H_2_O_2_. H_2_O_2_ is electrochemically active at the Pt/Ir electrode surface when a potential of +600 mV is applied, allowing for rapid (1 data point/sec or 1Hz) detection of changes in the concentration of extracellular glutamate in the brain. AscOx catalyzes the breakdown of ascorbate to dehydroascorbate and H_2_O, attenuating oxidation and detection of ascorbate at the electrode surface. NO-sensitive electrodes were generated using expired, heat inactivated GluOx electrodes. Insensitivity to glutamate was confirmed following 90-day room temperature incubation. Following room temperature incubation, electrodes were coated with a single layer of Nafion (5% wt/vol, Sigma) for 60 seconds and baked at 80°C for ten minutes. All biosensors interfaced with a PC via a wireless, Bluetooth® compatible, head-mounted potentiostat. A constant +600 mV electrochemical potential was applied to the electrode through the potentiostat during *in vitro* calibration as well as during *in vivo* recordings

### Antibodies and Viral vectors

In experiment 1, chicken anti-mCherry (RRID:AB_2716246, 1:2K, LS Biosciences) or chicken anti-GFP (RRID:AB_300798, 1:2K, Abcam) primary antisera were used for immunohistochemical detection of the mCherry tag on the hM3Dq receptor, or the eGFP reporter from the empty vector control group. Goat anti-chicken secondary antisera conjugated to Alexa 594 or 488 was used to amplify mCherry and eGFP, respectively. The chemogenetic vector AAV5-CaMKIIα-hM3Dq-mCherry and control reporter-only vector AAV5-CaMKIIα-eGFP (Titer: ∼2×10^12^ vg/ml, 0.75 µl/hemisphere) were purchased from Addgene (Cambridge, MA).

## Methods

### Animal subjects and surgery

Male Sprague-Dawley rats (*N*=47) were purchased from Charles Rivers Laboratories (Wilmington, MA), weighing 250-300 grams upon arrival. Rats were individually housed within a temperature and humidity-controlled room on a reverse light/dark cycle (lights off at 6 AM, lights on at 6 PM). Rats were acclimated to the vivarium for at least 3 days before undergoing surgery, during which rats had food and water available *ad libitum*. All animal use protocols were approved by the Institutional Animal Care and Use Committee of the Medical University of South Carolina (animal use protocol # 107172) and were performed according to the National Institutes of Health Guide for the Care and Use of Laboratory Animals (8^th^ ed., 2011). On the day of surgery, rats were anesthetized with an intraperitoneal ketamine (66 mg/kg) and xylazine (1.33 mg/kg) injection, and received ketorolac (2 mg/kg, i.p.) for analgesia. Rats in experiment 1 were secured in a stereotaxic apparatus and received a bilateral intra-PL cortical (AP: +2.8mm, ML: +/- 0.6mm, DV: −3.8 mm relative to bregma) microinjection of either a chemogenetic or reporter-only control vector (see materials). Rats in experiments 2 and 4 received a chronic indwelling jugular silastic catheter surgery as described previously (Siemsen *et al*. 2018a; Siemsen *et al*. 2018b; Siemsen *et al*. 2019), and received a single intravenous infusion of Cefazolin to prevent bacterial infection. All rats received a unilateral intra-NAcore guide cannula implantation following 4 weeks of virus expression (Experiment 1), immediately following catheterization (Experiment 2), in lieu of catheterization (Experiment 3) or 24 hours following the final cocaine SA session (Experiment 4, see below). Guide cannulae were unilaterally implanted ∼2 mm above the NAcore (+1.7mm AP, −1.6mm ML, −5mm DV relative to bregma). Following surgery, rats in Experiments 1-3 were returned to the vivarium for 5 days of post-operative recovery care. During this period, rats in Experiments 2-3 were handled daily and catheters were flushed daily with 50 µl of TCS (Access Technologies, Skokie, IL) to maintain catheter patency. Following guide cannulae implantation, rats in Experiment 4 underwent one week of forced homecage abstinence whereby food and water were available *ad libitum*. No specific randomization method was performed to allocate subjects in this study.

### *In vitro* calibrations of GluOx and NO-sensitive electrodes

Electrodes were screened prior to use for analyte sensitivity (nA/µM), selectivity (ratio of analyte:interferent nA/µM), and limit of detection according to Ferreira and colleagues (Ferreira *et al*. 2005). Briefly, sensitivity was calculated by dividing the difference in the nA response of each analyte by the prior baseline in nA. All responses for each analyte or interferent were then converted to nA/µM. Selectivity was then calculated by expressing electrode sensitivity of each analyte relative to that of the specific interferent. GluOx electrodes were calibrated using glutamate (8 µM) and response to the interferent ascorbic acid (100 µM) was also assessed. NO-sensitive electrodes were calibrated using the NO donor DETA (10 µM, yielding 100 nM NO). Potential responses from interferents including ascorbic acid (100 µM), glutamate (8 µM), dopamine (4 µM) were also analyzed. All solutions for *in vitro* calibrations were made in ddH20, except for DETA (see above). The nA response of each electrode during *in vivo* recordings was converted to ΔnM for glutamate and NO, as described previously (Isherwood *et al*. 2018). Briefly, the ΔnM glutamate or NO was calculated using the slope of the linear regression of each electrode’s *in vitro* calibration, relative to a 15-minute pre-session home-cage baseline.

### Novel context exposure

Rats in Experiment 3 (Glutamate *n*=5, NO *n*=5) received intra-NAcore guide cannula implantation and were allowed 5 days of recovery with daily handling as described above. One rat from the eGFP control group in Experiment 1 was used for a glutamate recording in Experiment 3 prior to the rats’ completion of the DREADD-evoked glutamate recording. On the last day of recovery, obturators were removed, and glutamate or NO-sensitive electrodes were lowered into the guide cannula, and rats were then returned to the vivarium overnight with the head-mounted potentiostat installed, however the potentiostat was not active during this period. The next morning the potentiostat was activated, the recordings began, and rats were placed in a dark room for 30-60 minutes until a steady baseline was achieved. Implantation of the electrode 15 hours prior to experimentation drastically improved the stability of the pre-experiment baseline response of the electrode. Rats were then exposed to a novel MedPC operant chamber with 2 interactable levers, a grated floor, and a house light for two hours where lever presses had no programmed consequence. Fluctuations in the nA response of glutamate- or NO-sensitive electrodes were recorded for two hours during the time when levers remained extended.

### Cocaine self-administration, extinction, and cue-induced reinstatement

In Experiment 4, rats were trained to either self-administer (SA) cocaine (*n*=14) or saline (*n*=11) for 10-12 days (2 hours/day) on a Fixed Ratio (FR) 1 schedule of reinforcement. Active lever presses elicited a light plus tone conditioned stimuli complex followed by a single infusion of cocaine hydrochloride (NIDA, Research Triangle Park, NC; 200 µg/50 µl bolus) followed by a 20 second timeout period. Inactive lever presses had no programmed consequence. After guide cannula implantation and recovery (described above), rats underwent extinction training until criterion was met (≤25 active lever presses over last 2 days of extinction). Following their final extinction session, obturators were removed from the guide cannula and either a glutamate or NO-sensitive electrode was lowered into the cannula; extending 3mm (including a 1mm sensing cavity) past the cannula tip. Animals were returned to the vivarium overnight with the head-mounted potentiostat installed. As described above for novelty, on the morning of reinstatement testing, rats remained in their home cage, placed in a dark room adjacent to the operant chambers (30-60 minutes) to collect baseline data. During cue-induced reinstatement, active lever presses elicited the light and tone conditioned cues, but no cocaine infusion, whereas inactive lever presses had no consequence. Active lever presses and cue presentations were time locked to the beginning and end of the recording session, and lever presses and cue presentations were extracted in 1-second bins through MedPC software.

### Transcardial perfusions, histology, and microscopy

Immediately following reinstatement testing, electrodes were extracted from the guide cannula. Rats were then heavily anesthetized with 1.5 ml of urethane (30% w/v, i.p.) and transcardially perfused with 120 ml 0.1M PBS (pH 7.4) followed by 180 ml of 5% formalin (60 ml/minute). Brains were extracted, post-fixed for 24 hours, and 60 µm coronal sections through the PL cortex and NAcore were collected using a vibrating microtome (Leica). Sections were mounted on glass slides and immediately cover slipped with Permount™ mounting medium to produce opaque tissue for probe localization (Fisher Scientific, Hampton, NH) using an inverted brightfield microscope (Evos). Disruption of brain tissue corresponding to the most ventral tip of the electrode was mapped according to the atlas of Paxinos and Watson. Animals were excluded if a significant (≥ 30%) portion of the 1mm sensing area was outside of the nucleus accumbens core. To verify the extent of virus expression, sections of the PL cortex and NAcore were immunohistochemically processed for either mCherry or eGFP as described previously (Giannotti *et al*. 2018). Briefly, 3-4 sections spanning the PL cortex and NAcore were blocked in 0.1M PBS containing 2% TritonX-100 and 2% normal goat serum (NGS) (PBST) for 2 hours at room temperature while shaking. Sections were then incubated in the appropriate primary antisera (see materials) in PBST overnight at 4°C while shaking. Sections were then washed 3×5 minutes in PBST and incubated in species-appropriate antisera (see materials) for two hours at room temperature while shaking. Sections were washed 3×5 minutes in PBST and then cover slipped with ProlongGold™ Antifade (Fisher Scientific). A Leica SP8 laser-scanning confocal microscope was used to image mCherry or eGFP expression in the PL cortex and NAcore.

### *In vivo* pharmacological validation of NO-sensitive electrodes

In experiment 2, drug-naïve rats (*n*=6) were infused i.v. with NaOH (3.2 µmol/320 µl) and responses of NO-sensitive electrodes were recorded for 30 minutes or until the nA response returned to the pre-experiment baseline (a minimum of 15 minutes), whichever occurred first. Rats were then infused i.v. with DETA (∼1.6 mg/kg, 3.2 µmol/320 ul) and NO-sensitive electrode responses were recorded in the same manner as vehicle. This protocol was performed in a counter-balanced manner, and each recorded epoch was normalized to a 5-15-minute baseline either before the experiment or prior to the second recorded epoch. A subset of rats (*n*=4) was then allowed an additional hour to reach a new, stable baseline. Rats were then infused i.v. with the vehicle for 7-NI (100% DMSO, 150 ul) followed by 7-NI (∼65.24 mg/kg, 60 µmol/150 µl). Vehicle was recorded for one hour. Given the variability in the time at which 7-NI effected the nA baseline response of NO-sensitive electrodes, we recorded NO-sensitive electrode responses for either one hour or until the response returned to the previous baseline (a minimum of 15 minutes). Each recorded epoch was normalized to a 10-15-minute baseline either before the experiment or before the second recorded epoch.

### Experimental design

All experiments were conducted in the morning; approximately 15 hours after electrode implantation. A schematic of the timeline for each experiment is shown in Figure 1. Experiment 1 was designed to validate *in vivo* detection of glutamate with chemogenetic-mediated activation of PL cortical excitatory neurons on the induction of NAcore glutamate. Experiment 2 was designed to validate *in vivo* detection of NO with intravenous infusions of DETA or 7-NI to increase or decrease NO concentrations in the NAcore *in vivo*, respectively. Experiment 3 and 4 were designed to quantify and compare glutamate and NO release in the NAcore of drug-naïve rats during exposure to a novel context or during cue-induced reinstatement of cocaine or saline seeking, respectively.

**Figure 1.**
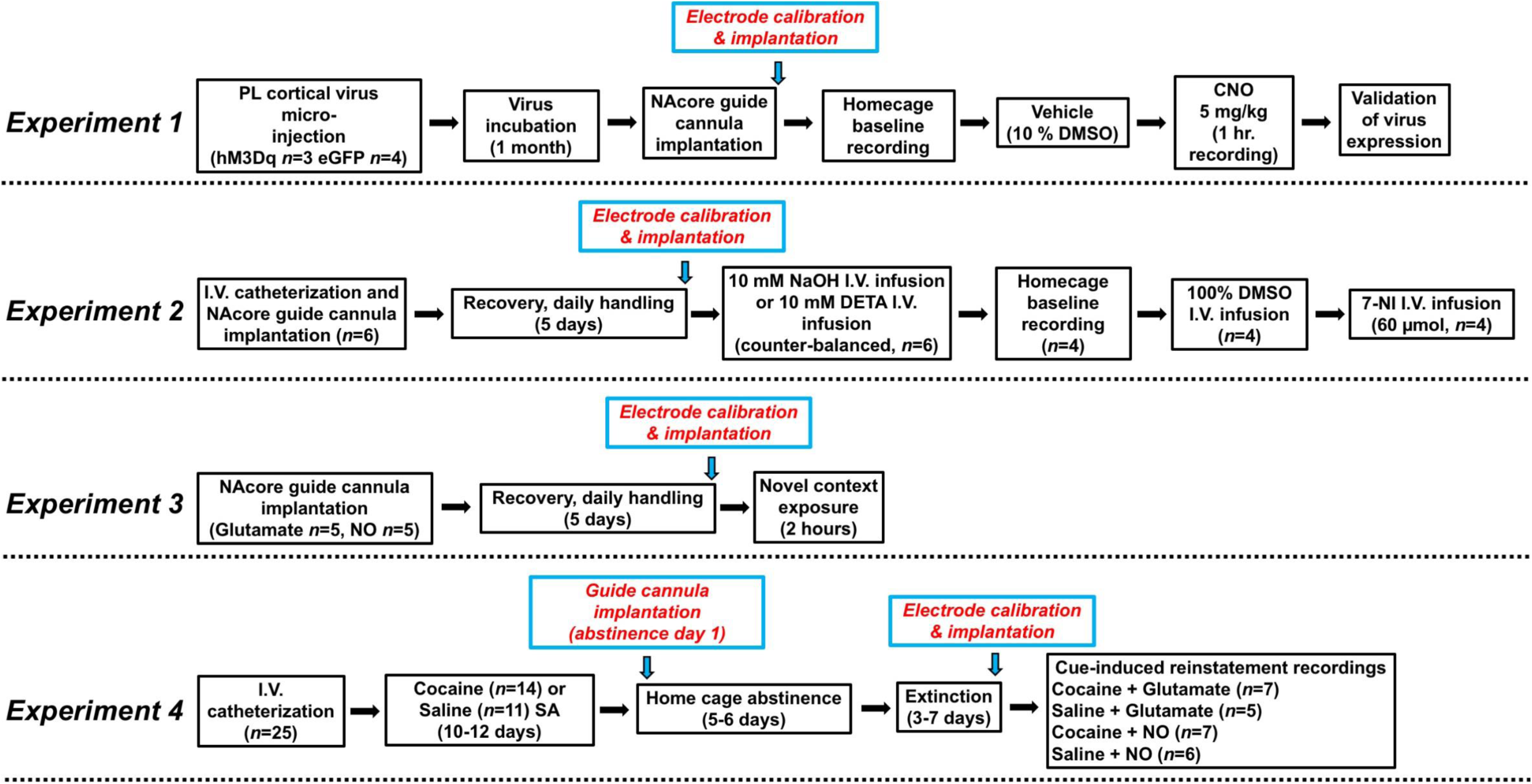
Experimental design for experiments 1-4.

### Statistical analyses

Sample sizes were estimated from previously published amperometric studies using identical GluOx biosensor recordings (Gass *et al*. 2011; Wakabayashi & Kiyatkin 2012). All data analysis was performed by an experimenter who was unblind to experimental groups. Amperometric data was analyzed with Sirenia® Acquisition software (v. 2.0.3, Pinnacle Technology). All statistical data was analyzed using GraphPad (Prism, version 8.1). Area Under the Curve (AUC) analyses were compared within groups or between multiple groups using a two-tailed paired t-test or one-way ANOVA with Tukey’s multiple comparison test when a significant main effect was observed, respectively. Reinstatement behavioral data was analyzed with a two-way repeated measures ANOVA with treatment (cocaine versus saline) as a between-subjects variable and time (extinction versus reinstatement) as a within-subjects variable followed by Bonferroni-corrected multiple comparison test when a significant interaction was revealed. 1 Hz measurements of glutamate and NO during reinstatement and novel context exposure were averaged in 5-minute bins (first hour of session) or 1-minute bins (first 15 minutes of session). A two-way repeated measures ANOVA was performed followed by Bonferroni-corrected pairwise comparison tests to compare across treatments within bins when a significant treatment by time interaction was revealed. A one-way repeated measures ANOVA was used when analyzing evoked responses across time within a single group, followed by Dunnett’s multiple comparison test to compare each binned data point to the baseline. *In vitro* calibrations were analyzed with linear regression. In experiments 3-4, the AUC for the entire two-hour session was used to detect statistical outliers according to Grubbs’ test (α=0.05) and these were removed from all analyses (*n*=2, see results). Data is expressed as the average +/- the standard error of the mean (SEM), and significance was set at *p*<0.05.

## Results

### *In vitro* validation of GluOx and NO-sensitive electrode

Figure 2A,B shows the surface chemistry of GluOx electrodes and a representative GluOx electrode calibration in response to ascorbic acid and glutamate, respectively. GluOx electrodes displayed a highly linear response to repeated glutamate additions (R^2^=0.997 +/- 0.001, Figure 2C). The sensitivity of GluOx electrodes for ascorbic acid and glutamate, the selectivity of GluOx electrodes for glutamate relative to ascorbic acid, and the limit of detection (LOD) for glutamate can be found in Table 1. These data are largely consistent with previously reported findings from independent laboratories using the same biosensors (Wakabayashi & Kiyatkin 2012).

**Table 1.**
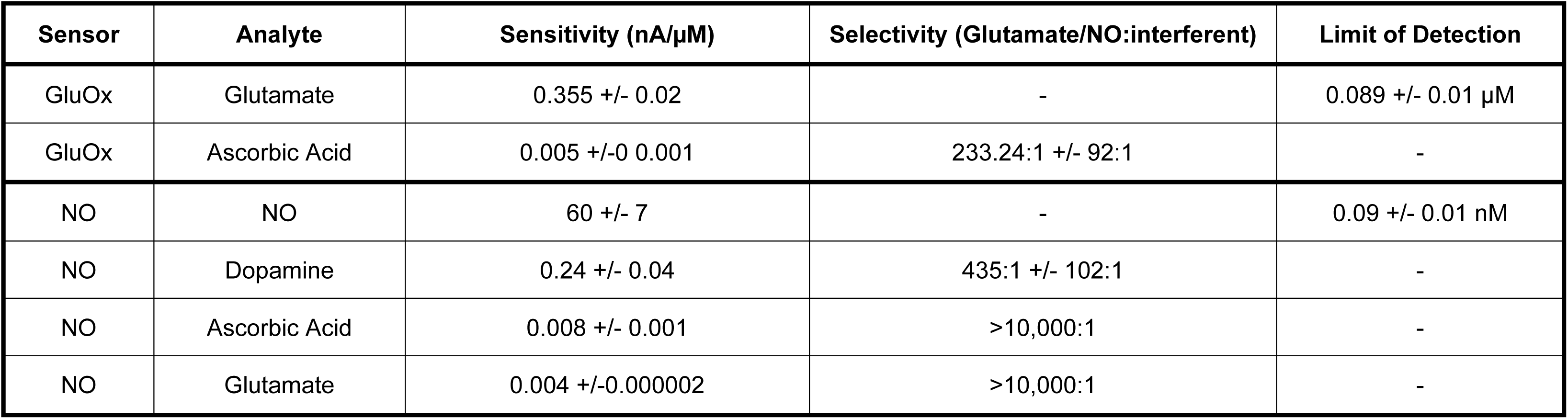
Sensitivity and selectivity calculations for all glutamate and NO electrodes used in experiments 1-4.

| Sensor | Analyte | Sensitivity (nA/μM) | Selectivity (Glutamate/NO:interferent) | Limit of Detection |
| --- | --- | --- | --- | --- |
| GluOx | Glutamate | 0.355 ± 0.02 | - | 0.089 ± 0.01 μM |
| GluOx | Ascorbic Acid | 0.005 ± 0.001 | 233.24:1 ± 92:1 | - |
| NO | NO | 60 ± 7 | - | 0.09 ± 0.01 nM |
| NO | Dopamine | 0.24 ± 0.04 | 435:1 ± 102:1 | - |
| NO | Ascorbic Acid | 0.008 ± 0.001 | >10,000:1 | - |
| NO | Glutamate | 0.004 ± 0.000002 | >10,000:1 | - |

**Figure 2.**
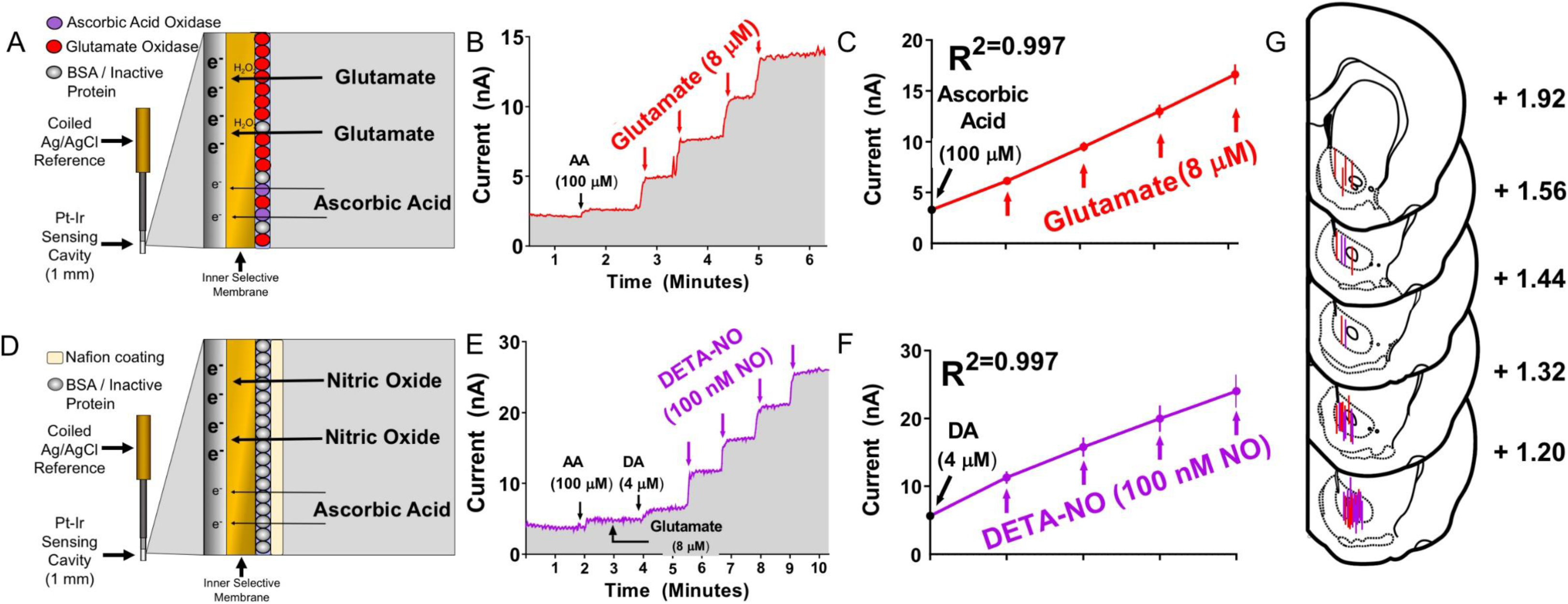
GluOx and NO-sensitive electrodes’ surface chemistry, *in vitro* calibrations, and histological validation. **A**. Schematic illustrating the surface chemistry used to detect glutamate via GluOx. **B**. Representative *in vitro* GluOx biosensor calibration. Note the low sensitivity to ascorbic acid (100 µM) relative to repeated glutamate (8 µM) additions. **C**. Quantification of the linearity of all GluOx electrodes (*N*=21) used for all experiments. **D**. Schematic illustrating the surface chemistry used to detect NO. **E**. Representative *in vitro* NO-sensitive electrode calibration. Note that NO-sensitive electrodes show little sensitivity to ascorbic acid (100 µM), near-zero sensitivity to glutamate (8 µM), and low sensitivity to dopamine (4 µM) relative to repeated DETA-NO (10 µM DETA converted to 100 nM NO) additions. **F**. Quantification of the linearity of all NO-sensitive electrodes (*N*=19) used for all experiments. **G**. Histological verification of the most ventral tip for all glutamate (red) and NO (purple) electrodes. Note most electrode placements are confined to the caudal portion of the NAcore.

Figure 2D shows a schematic of the NO-sensitive electrode design. Figure 2E shows a representative NO-sensitive electrode calibration in response to ascorbic acid, glutamate, dopamine, and DETA. NO-sensitive electrodes displayed a highly linear response to additions of DETA (R^2^=0.997 +/- 0.0008, Figure 2F). The sensitivity of NO-sensitive electrodes to ascorbic acid, glutamate, dopamine, and NO, their selectivity for NO relative to multiple interferents, and the LOD for NO can be found in Table 1. NO-sensitive electrodes were rendered 99% insensitive to glutamate relative to GluOX electrodes, with average nA/μM glutamate responses below what was observed for ascorbic acid in either configuration. Figure 2G shows the anatomical mapping of the 1mm sensing area of each electrode. The majority of electrode placements are localized to the caudal part of the NAcore by design, as this sub-region is recruited by cues predicting appetitive stimuli (Hamel *et al*. 2017).

#### Experiment 1: *In vivo* validation of Glutamate-sensitive electrodes

To validate the utility of GluOx electrodes in detecting evoked glutamate in awake and freely moving rats, we transduced PL cortical glutamatergic neurons with excitatory (hM3Dq) DREADDs. These neurons were selected as they both project to the NAcore (Gabbott *et al*. 2005), and are a likely source of the cue-induced glutamate release in the NAcore that drives relapse (Stefanik *et al*. 2016). Figure 3A shows a schematic depicting the surgical design and probe placement. Representative micrographs of AAV5-CaMKIIα-hM3Dq-mCherry (*n*=3) and AAV5-CaMKIIα-eGFP (*n*=4) expression in the NAcore (top) and PL cortex (bottom) are shown in the left and right columns, respectively, in Figure 3B. Figure 3C shows the average change in glutamate in eGFP- and hM3Dq-expressing rats. Figure 3D shows quantification of data in Figure 3C. A two-way repeated-measures ANOVA indicated a significant virus by drug interaction (F(1,5)=126.5, *p*<0.0001). Bonferroni-corrected pairwise comparison test indicated that hM3Dq-expressing rats showed significantly greater AUC (baseline=0 nM) following CNO injections than eGFP-expressing controls (*p*<0.001), as well as a significantly greater AUC compared to its own vehicle (*p*<0.0001), whereas there was no difference between groups in the response to vehicle (*p*=0.52).

**Figure 3.**
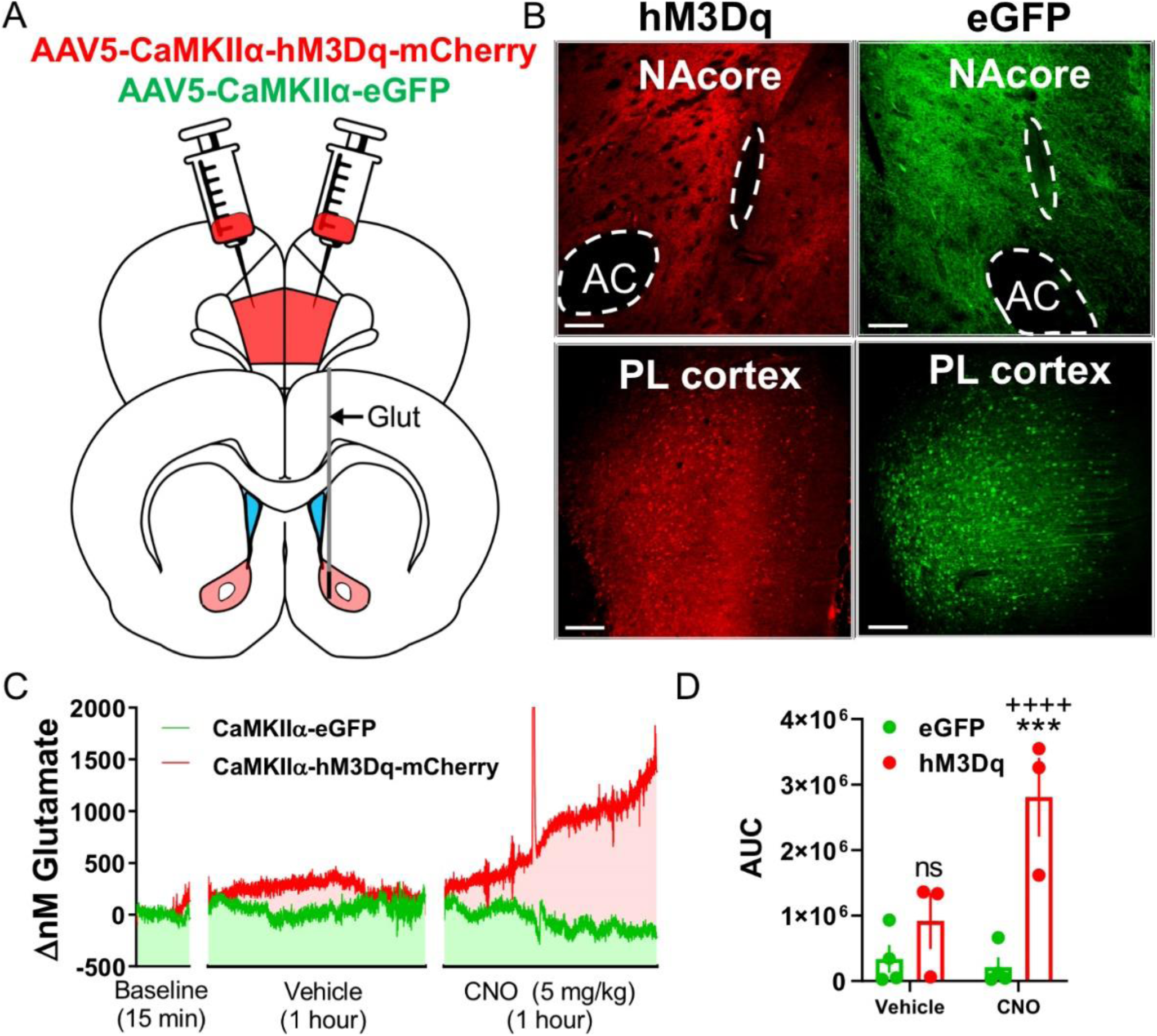
Chemogenetic-evoked detection of glutamate in the NAcore of freely moving rats. **A**. Schematic illustrating virus microinjection site (PL cortex) and recording site (NAcore). **B**. Representative terminal expression in the NAcore of mCherry (hM3Dq) or eGFP (empty virus control) expression within the proximity of the probe within the NAcore (top) and representative virus expression from each group in the PL cortex (bottom). **C**. Average change in glutamate concentration (1Hz resolution) within the NAcore following vehicle (10% DMSO) and CNO (5 mg/kg, i.p.) in hM3Dq (*n*=3, red) and eGFP (*n*=4, green) rats relative to a 15-minute homecage baseline. **D**. Quantification of the area under the curve (AUC, baseline = 0 nM) of the change in glutamate in the NAcore following vehicle and CNO injections. Two-way repeated-measures ANOVA with Bonferroni-corrected pairwise comparison test, \*\*\**p*<0.001 compared to eGFP, ++++*p*<0.0001 compared to vehicle. Scale bars=200 µM.

#### Experiment 2: *In vivo* validation of NO-sensitive electrodes

To validate the utility of NO electrodes in detecting changes in NO levels in awake and freely moving rats we utilized the same NO donor used in our calibrations as well as an nNOS inhibitor. In experiment 2, drug-naïve rats (*n*=6) were i.v. catheterized and implanted with unilateral guide cannula in the NAcore as described above. Rats then received i.v. infusions of the vehicle for DETA (10mM NaOH) followed by a single i.v. infusion of DETA while recording responses of NO-sensitive electrodes in a counter-balanced manner. A representative trace is shown in Figure 4A. A two-tailed paired t-test indicated a significant effect of DETA compared to vehicle when analyzing the AUC (baseline=0 nM) of each epoch (t(5)=2.84, *p*<0.05, Figure 4B). We also calculated the maximum peak height achieved following either vehicle or DETA treatment. A two-tailed paired t-test indicated that DETA infusions evoked a significantly greater peak nM NO response when compared to vehicle infusions (t(5)=7.56, *p*<0.001, Figure 4C).

**Figure 4.**
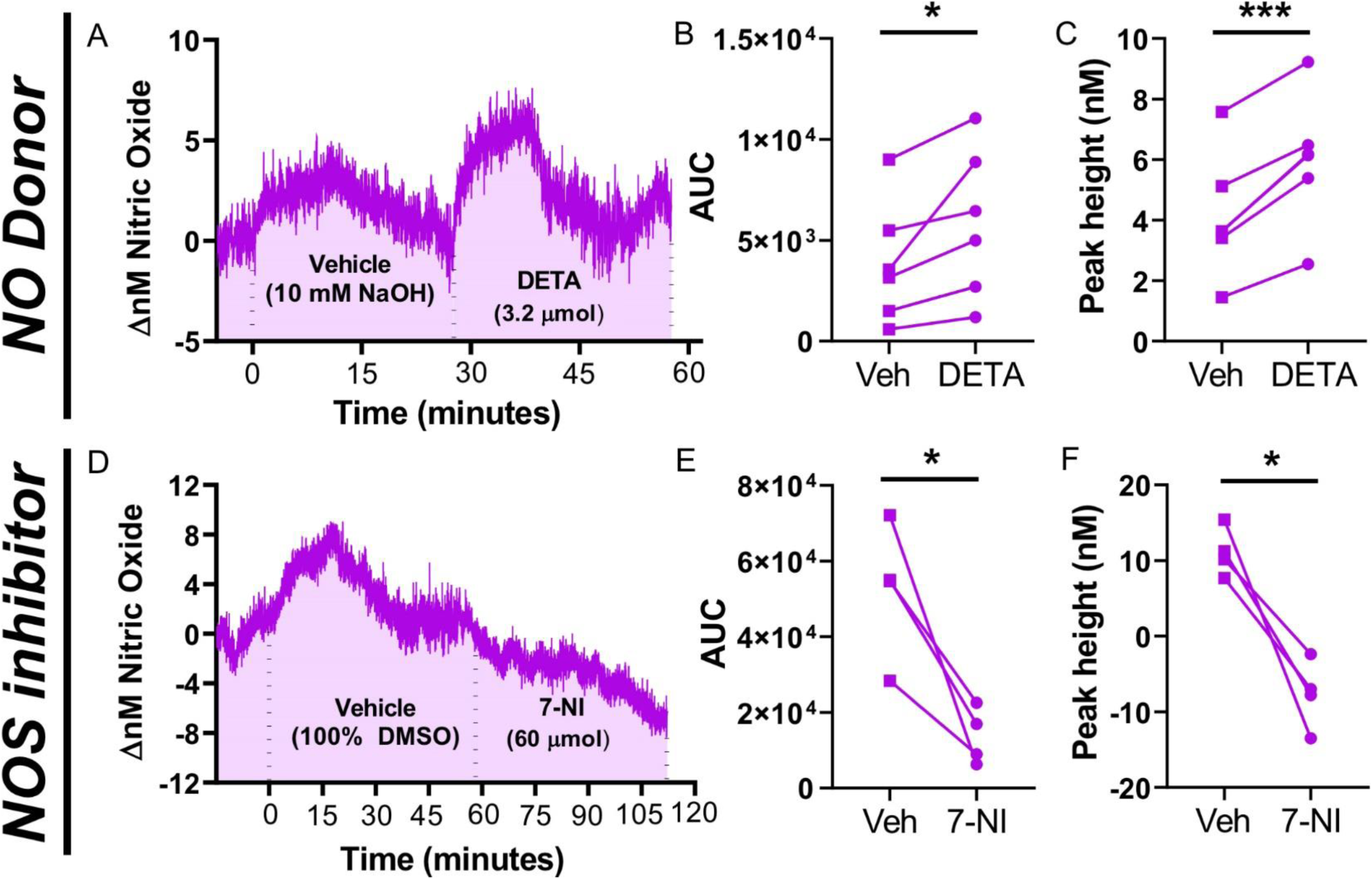
*In vivo* intravenous pharmacological validation of NO-sensitive electrodes in freely moving rats. **A**. Representative trace showing the change in the concentration of NO in the NAcore measured once per second in response to i.v. infusions of vehicle (10 mM NaOH, *n*=6) and DETA (3.2 µmol, *n*=6). **B**. Quantification of the AUC (baseline = 0 nM) and **C**. peak height (in nM) in response to vehicle and DETA for all replicates. **D**. Representative trace showing the change in NAcore NO concentrations measured once per second in response to i.v. infusions of vehicle (100 % DMSO, *n*=4) and 7-NI (60 µmol, *n*=4). **E**. Quantification of the AUC (baseline = −10 nM) and **F**. peak height (in nM) in response to vehicle and 7-NI. B, C, E, F: two-tailed paired t-test, \**p*<0.05, \*\*\**p*<0.001 compared to respective vehicle.

A subset of rats (*n*=4) was returned then to their homecage for one hour and given an i.v. infusion of the vehicle for 7-NI (DMSO) followed by an i.v. infusion of 7-NI while recording responses of NO-sensitive electrodes. An individual representative trace is shown in Figure 4D. A two-tailed paired t-test indicated that 7-NI significantly decreased the signal of NO-sensitive electrodes relative to vehicle; observed as a decrease in the baseline values when analyzing the AUC (baseline=-10 nM) of each epoch (t(3)=3.97, *p*<0.05, Figure 4E). 7-NI also showed a significantly lower peak height compared to vehicle (t(3)=5.17, *p*<0.05, Figure 4F).

#### Experiment 3. Exposure to a novel environment increases glutamate and NO in the NAcore

Given that exposure to a novel context has previously been shown to increase glutamate (Saulskaya & Marsden 1995) and co-products of NO synthesis in the NAcore (Saul’skaya & Sudorgina 2014), we first set out to quantify the release of glutamate (*n*=5) and NO (*n*=5) produced by exposing rats to a novel context. One rat undergoing NO recordings during novel context exposure was removed from the analysis as a statistical outlier. While there were no programmed consequences for responding on either lever, rats undergoing glutamate and NO recordings pressed the levers an average of 15+/-6 and 18+/-9 times, respectively. Figure 5A shows the mean change in glutamate in the NAcore during two hours of exposure to the novel context. Figure 5B shows binned data (5-minute bins) for the mean change in glutamate in the NAcore during the first hour of exposure to the novel context relative to a pre-session home-cage 15-minute baseline. A repeated-measures one-way ANOVA revealed a significant effect of time (F(12,48)=10.36, *p*<0.0001), and Dunnett’s multiple comparison test comparing each bin to the baseline revealed that exposure to a novel context engages significant glutamate release in the NAcore within 10 minutes (*p*<0.001). As our sampling rate is 1 Hz, we focused on the first 15 minutes of the two-hour session to increase the temporal precision of our analyses, and to gain a better understanding of the precise time (in minutes) at which glutamate or NO is increased in the NAcore upon exposure to a novel context. When the first 15 minutes of novel context exposure was expressed in 1-minute bins, a repeated-measures one-way ANOVA revealed a significant effect of time (F(15,60)=30.52, *p*<0.0001). Dunnett’s multiple comparison test indicated that exposure to a novel context takes 2 minutes to evoke significant glutamate release in the NAcore (*p*<0.01, Figure 5F).

**Figure 5.**
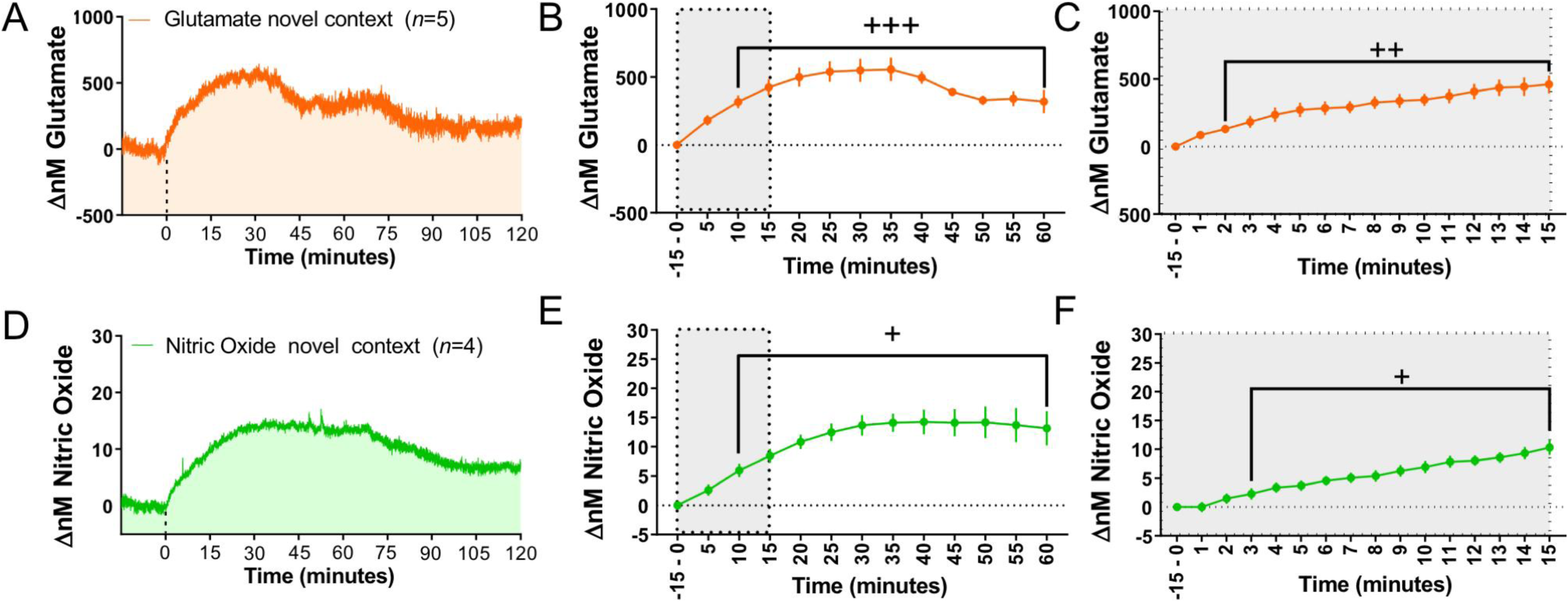
Exposure to a novel context with interactable levers increases glutamate and NO in the NAcore. **A**. Average change in glutamate in the NAcore during exposure to a novel context (*n*=5). **B**. Raw data shown in A expressed in 5-minute bins over the first hour of the session relative to a 15-minute baseline. **C**. One-minute binned data over the first 15 minutes of novel context exposure. **D**. Average change in NO in the NAcore during exposure to a novel context (*n*=4). **E**. Raw data shown in D expressed in 5-minute bins over the first hour of context exposure relative to a pre-session baseline. **F**. Raw data shown in D expressed in 1-minute bins during the first 15 minutes of novel context exposure. B, C, E, F: One-way repeated-measures ANOVA with Dunnett’s multiple comparison test, +*p*<0.05, ++*p*<0.01, +++*p*<0.001 compared to baseline. Grey rectangles in B,E indicate the epoch whereby analysis was performed in C,F.

We also performed the same analysis for NO. The mean change in the concentration of NO in the NAcore during exposure to a novel context is shown in Figure 5D. When data was expressed in 5-minute bins, a repeated-measures one-way ANOVA revealed a significant main effect of time (F(12,36)=11.71, *p*<0.0001). Akin to glutamate, NO is increased in the NAcore within 10 during exposure to a novel context, according to Dunnett’s multiple comparison test (*p*<0.05, Figure 5E). When the first 15 minutes of novel context exposure was analyzed in 1-minte bins, a repeated-measures one-way ANOVA revealed a significant main effect of time (F(15,45)=21.61, *p*<0.0001). In contrast to glutamate, Dunnett’s multiple comparison test indicated that novel context exposure increased NO in the NAcore within 3 minutes (*p*<0.05, Figure 5F).

#### Experiment 4. Cue-induced reinstatement increases glutamate and NO in the NAcore

Of the 25 rats that were included in the beginning of the experiment, 6 rats were removed for either lack of reinstatement (Cocaine SA-glutamate recordings *n*=2), cannula placement outside of the NAcore (Cocaine SA-NO recordings *n*=2, Saline SA-NO recordings *n*=1), or was a statistical outlier (Cocaine SA-NO recordings *n*=1). Figure 6A shows SA and extinction behavior for both cocaine and saline SA animals undergoing either glutamate or NO recordings. Saline and cocaine SA animals received an average of 4.7+/-0.45 and 5.2+/-0.63 days of extinction, respectively. Figure 6B shows cue-induced reinstatement behavior for all SA rats, used in either glutamate or NO recordings. A two-way repeated-measures ANOVA revealed a significant treatment by time interaction when comparing extinction and cue-induced reinstatement in cocaine compared to saline SA rats (F(1,17)=53.32, *p*<0.0001). Bonferroni-corrected pairwise comparison tests indicated that cocaine SA rats showed greater levels of responding on the active lever averaged across the last two days of extinction (*p*<0.05) and significantly higher presses on the formerly active lever during reinstatement compared to saline SA rats (*p*<0.0001). Figure 6C-D shows a heat map of the average responses on the active lever during cue-induced reinstatement in saline and cocaine SA rats in 1-minute bins over the full two-hour session or first 15 minutes, respectively.

**Figure 6.**
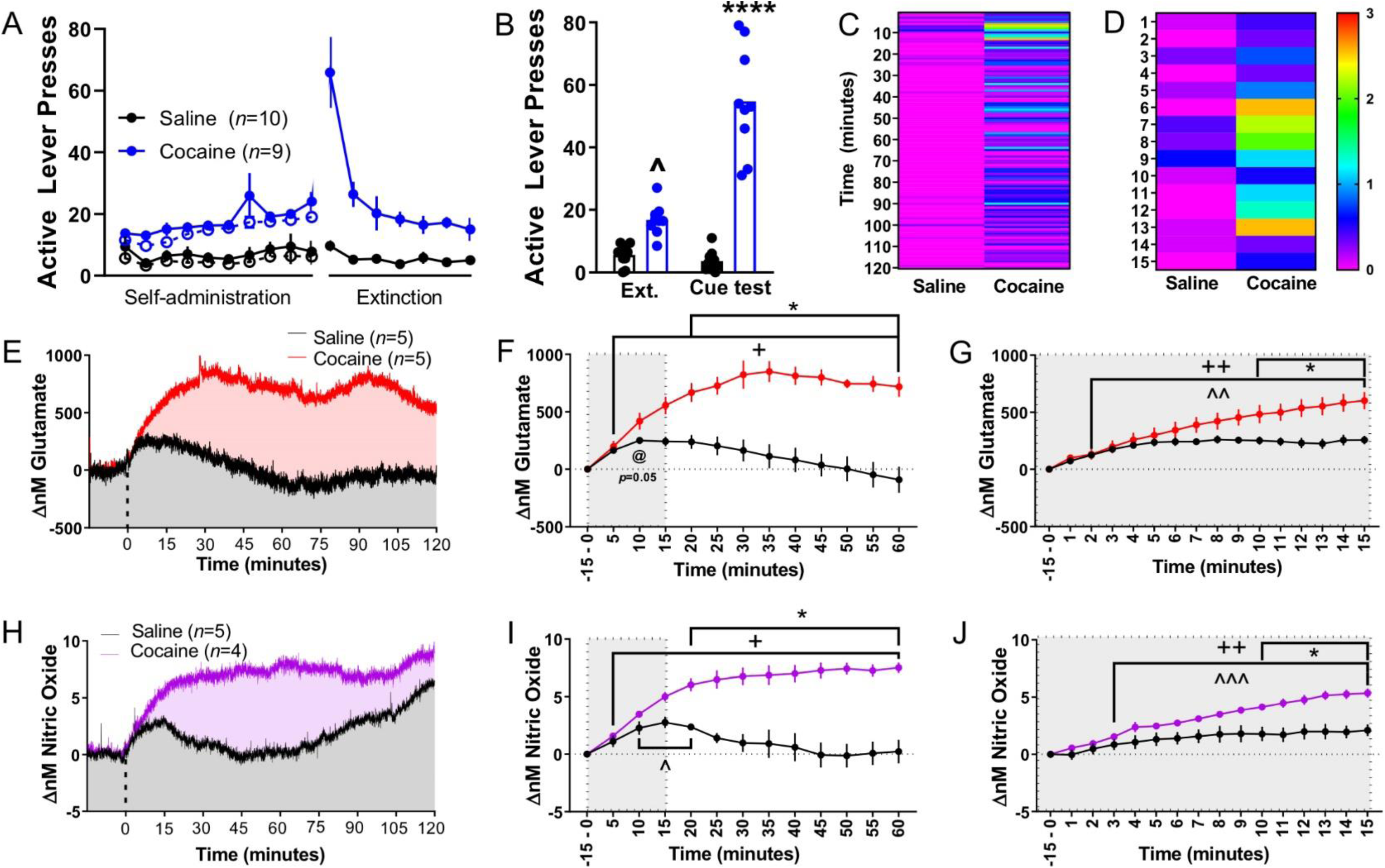
Cue-induced reinstatement increases glutamate and NO in the NAcore. **A**. Active lever presses in saline (black, *n*=10) and cocaine (blue, *n*=9) SA rats during self-administration and extinction. Closed circles indicate active lever presses over the last 10 days of self-administration and 7 days of extinction. Open circles indicate infusions earned. **B**. Average active lever presses over the last 2 days of extinction (left) and during cue-induced reinstatement in saline and cocaine SA rats. **C-D**. Heat map of the average active lever presses in one-minute bins for the full two-hour session (C) as well as during the first 15 minutes of reinstatement (D) in saline and cocaine SA rats. Warmer colors indicate a greater number of lever presses, whereas cooler colors indicate low lever pressing. **E**. Average change in glutamate in the NAcore of saline (black, *n*=5) and cocaine (red, *n*=5) SA rats during cue-induced reinstatement relative to a 15-minute pre-session baseline. **F**. Five-minute binned data of the raw data shown in E over the first hour of reinstatement. **G**. One-minute binned data of the raw data shown in E over the first 15 minutes of reinstatement. **H**. Average change in NO in the NAcore of saline (black, *n*=5) and cocaine (purple, *n*=4) SA rats. **I**. Five-minute binned data of the raw data shown in H. **J**. One-minute binned data of the raw data shown in H over the first 15 minutes of reinstatement. B: Two-way repeated-measures ANOVA with Bonferroni-corrected pairwise comparison tests. ^*p*<0.05 compared to saline extinction, \*\*\*\**p*<0.0001 compared to saline reinstatement. F, G, I, J: two-way repeated measures ANOVA with Bonferroni corrected pairwise comparison test \**p*<0.05 compared to saline. F, G, I, J: One-way repeated-measures ANOVA with Dunnett’s multiple comparison test +*p*<0.05, ++*p*<0.01, compared to baseline (cocaine), ^*p*<0.05, ^^*p*<0.01, ^^^*p*<0.001 compared to baseline (saline), @*p*=0.05 compared to baseline (saline).

Following extinction rats (Cocaine *n*=5, Saline *n*=5) were implanted with GluOx electrodes and underwent cue-induced reinstatement the following day. Figure 6E shows the mean change in the concentration of glutamate during cue-induced reinstatement in cocaine and saline SA rats, relative to a 15-minute homecage baseline. We then compared the AUC of the full two-hour session from these groups to the AUC of the glutamate response from novel context exposure, and a one-way ANOVA revealed a significant main effect of treatment (F(2,12)=21.77, *p*<0.0001, Figure S1A). Tukey’s multiple comparison test indicated that cocaine SA animals showed a significantly greater AUC than both saline SA (*p*<0.0001) and novel context exposed rats (*p*<0.01), yet saline SA was not significantly different than novel context exposure. When analyzing the first hour of reinstatement in 5-minute bins in cocaine and saline SA rats, a two-way repeated-measures ANOVA indicated a significant treatment by time interaction (F(12,96)=15.18, *p*<0.0001, Figure 6F). Bonferroni-corrected pairwise comparison test indicated that cocaine SA rats showed a significantly larger change in glutamate 20 minutes into reinstatement (*p*<0.05) compared to saline SA rats. In order to compare each bin to the baseline, a one-way repeated-measures ANOVA was performed for each group. For both cocaine (F(12,48)=33.75, *p*<0.001) and saline (F(12,48)=3.59, *p*<0.001), this analysis revealed a significant main effect of time. Dunnett’s multiple comparison test indicated that cue-induced reinstatement leads to an increase in NAcore glutamate within 5 minutes (*p*<0.05) for cocaine SA animals, yet there was no significant timepoint whereby glutamate was increased compared to baseline, although the 10-minute bin showed marginal significance (*p*=0.05), in saline SA animals.

As described above for novel context exposure, we next focused on the first 15 minutes of reinstatement. This was done both because this epoch has been shown to be particularly relevant for the glutamate-induced plasticity within the NAcore mediating cue-induced reinstatement (Gipson *et al*. 2013). Figure 6G shows the first 15 minutes analyzed in 1-minute bins. A two-way repeated-measures ANOVA revealed a significant treatment by time interaction (F(15,120)=13.07, *p*<0.0001). Interestingly, Bonferroni-corrected pairwise comparison tests indicated that cocaine and saline SA rats do not show a discernible difference in glutamate release until 10 minutes into reinstatement (*p*<0.05). To compare each bin to the home-cage baseline within each group, a one-way repeated-measures ANOVA was performed. The results revealed a significant main effect of time for both cocaine (F(15,60)=50.78, *p*<0.0001) and saline (F(15,60)=10.33, *p*<0.0001) SA rats. Dunnett’s multiple comparison test indicated that cue-induced reinstatement significantly increased NAcore glutamate levels within 2 minutes in both saline and cocaine SA rats, relative to a 15-minute homecage baseline (*p*<0.01).

A subset of rats (Cocaine *n*=4, Saline *n*=5), were implanted with NO-sensitive electrodes and then underwent cue-induced reinstatement of cocaine seeking the next day as described above. Figure 6H shows the average change in the concentration of NO in the NAcore during the two-hour reinstatement session. When we analyzed the AUC of NO recordings in cocaine and saline SA rats along with the NO response from novel context-exposed animals in Experiment 3, a one-way ANOVA revealed a significant main effect of treatment (F(2,10)=26.81, *p*<0.0001, Figure S1B). Tukey’s multiple comparison test indicated that both cocaine (*p*<0.01) and novel context exposed animals (*p*<0.0001) showed significantly higher evoked NO levels when compared to saline SA animals. However, novel context-exposed rats displayed elevated levels of NO release relative to cocaine SA animals during cued reinstatement (*p*<0.05). Figure 6I shows the first hour of reinstatement in 5-minute bins for cocaine and saline SA rats undergoing cue-induced reinstatement. A two-way repeated-measures ANOVA revealed a significant treatment by time interaction (F(12,84)=17.99, *p*<0.0001). Bonferroni-corrected pairwise comparison tests indicated a significant difference between cocaine and saline SA rats by 20 minutes of cue-induced reinstatement (*p*<0.05). To compare each bin to the home-cage baseline within each group, a one-way repeated-measures ANOVA was performed which revealed a significant main effect of time for both cocaine (F(12,36)=56.57, *p*<0.0001) and saline (F(12,48)=3.59, *p*<0.001) SA rats. Dunnett’s multiple comparison test indicated that cocaine and saline SA rats show cue reinstatement-induced increases in NO in the NAcore within 5 and 10 minutes, respectively. This increase was sustained for the remainder of reinstatement in cocaine SA rats, but returned to baseline levels by 25 minutes in saline SA rats.

We also analyzed the first 15 minutes of reinstatement in rats undergoing NO recordings in 1-minute bins. A two-way repeated measures ANOVA revealed a significant treatment by time interaction when analyzing the first 15 minutes in 1-minute bins (F(15,105)=9.91, *p*<0.0001, Figure 6J). Akin to glutamate, Bonferroni-corrected pairwise comparison test indicated a significant difference between cocaine and saline SA rats by 10 minutes into reinstatement (*p*<0.05). A repeated-measures one-way ANOVA revealed a significant main effect of time in both cocaine (F(15,45)=51.78, *p*<0.0001) and saline (F(15,60)=20.89, *p*<0.0001) SA rats. Dunnett’s multiple comparison test indicated that both saline and cocaine SA animals displayed increased cue reinstatement-induced NO in the NAcore by 3 minutes.

## Discussion

### Summary of findings

The first primary finding of this study is that cue-induced reinstatement of cocaine seeking elevates both glutamate and NO in the NAcore in a time-dependent manner. Dysregulated glutamate transmission in the NAcore is implicated in cue-induced seeking for multiple classes of drugs of abuse (Scofield *et al*. 2016a). Recently, a link between cue-induced glutamate release, activation of nNOS interneurons, and the induction of NO release in the NAcore has been made (Smith *et al*. 2017). Here we provide quantitative analysis, for the first time, of the extent of NAcore NO release during cue-induced reinstatement of cocaine seeking. Taken together, these data add to the growing body of literature indicating a critical role of NO signaling in the striatum in regulating behavioral responses to psychostimulants (Itzhak *et al*. 1998a; Itzhak *et al*. 1998b). The second primary finding of this study is that induction of a glutamate response in the NAcore precedes the NO response regardless of the context which evokes a response, even despite differences observed in the magnitude of glutamate and NO release during cue-induced reinstatement and exposure to a novel context. Taken together, these data contribute to better overall understanding of the relationship between evoked glutamate and NO release in the NAcore.

### *In vivo* validation of Glutamate- and NO-sensitive electrodes

Given that we are the first to perform amperometry-based freely moving measurements of cocaine cued reinstatement-evoked glutamate and NO release, we characterized not only the electrodes sensitivity and selectivity *in vitro*, but also performed positive control experiments to demonstrate the ability of these electrodes to detect glutamate and NO in awake and freely moving animals. The results of our chemogenetic experiments reveal that we were readily able to detect elevated glutamate following activation of glutamatergic cortical neurons with excitatory DREADDs. These data demonstrate that the GluOx electrodes readily detect glutamate release in the NAcore in freely moving rats, following DREADD-based stimulation of cortical glutamatergic projection neurons directly linked to cued reinstatement of cocaine seeking (Stefanik *et al*. 2016).

The results of our pharmacological experiments reveal that i.v. infusions of the NO donor DETA-NONOate or the relatively selective nNOS inhibitor, 7-NI, can increase or decrease brain NO levels, respectively. These data demonstrate that a large portion of the signal detected by our NO-sensitive electrodes is indeed NO. A previous study found that chronic (four i.v. bolus doses separated by 15 minutes followed by 6 daily i.p. injections) treatments with DETA at 0.4 mg/kg significantly increases cGMP (production of which is enhanced following activation of the NO receptor soluble guanylyl cyclase (sGC)) in the cortex (Zhang *et al*. 2001). Taken together with our results, these data indicate that i.v. administration of DETA increases brain NO levels. We observed that i.v. infusions of the same volume of the vehicle for DETA (10mM NaOH) increased the signal of NO-sensitive electrodes, albeit significantly lower than what was observed following administration of DETA. Given that administration of DETA produced a distinct kinetic profile (i.e. transient elevation) relative to the vehicle, and that the AUC and peak height were both significantly greater, these data demonstrate that NO-sensitive electrodes can detect increases in NO levels in the NAcore.

Pharmacological inhibition of nNOS has historically been the gold standard for elucidating the role of nNOS-dependent NO production, and 7-NI was selected in our experiments as it possesses greater selectivity for nNOS relative to other NOS isoforms (Babbedge *et al*. 1993; Moore *et al*. 1993). Further, pretreatment of rats with systemic injections of 7-NI prior to daily cocaine injections suppresses cocaine conditioned place preference, an effect absent in nNOS^-/-^mice, indicating both that nNOS activation is implicated in the maladaptive learning associated with the development and expression of CPP and that 7-NI is capable of suppression of nNOS activity (Itzhak *et al*. 1998b; Atalla & Kuschinsky 2006). As 7-NI is insoluble in water, we used DMSO as vehicle in our i.v. studies. We found that DMSO transiently increased NO release and that administration of 7-NI (∼65 mg/kg, 60 µmol/150µl) reverted this increase and also dropped the signal below the pre-DMSO baseline in the NAcore. DMSO itself increases vasodilation and elevates cGMP, a downstream effector of NO, in the rat aorta through a eNOS-dependent manner, providing evidence for the DMSO-induced NO release we observed in our studies (Kaneda *et al*. 2016).Taken together, these data demonstrate that NO-selective electrodes reliably detect both increases and decreases in NO concentrations in awake animals.

### Exposure to a novel context increases glutamate and NO release in the NAcore

Several studies describe enhanced glutamate release in the NAcore during exposure to a novel context. As an example, exposing rats to a novel fear-conditioning chamber, in the absence of tone or shock delivery, elevates extracellular glutamate in the NAcore, which returns to baseline upon re-exposure of an animal to their home-cage (Saulskaya & Marsden 1995). Moreover, exposing mice to a novel, complex environment (a novel context with interactable objects), increases Fos protein expression in the NAcore, likely as a result of enhanced glutamate and dopamine release (Rinaldi *et al*. 2010). Finally, biosensor recordings using the same GluOx electrodes used here indicate a transient increase in the current response of GluOx electrodes implanted in the NAcore during exposure to a naturally, arousing stimulus (brief tail pinch) (Wakabayashi & Kiyatkin 2012). We show here that exposure to a novel SA chamber with interactable levers elicits an increase of ∼500 nM of glutamate in the NAcore, which is evident within 2 minutes of exposure to the novel context and is sustained for at least the first hour of the exposure. These data are in agreement with what has been reported by others (Saulskaya & Marsden 1995; Rinaldi *et al*. 2010) and demonstrates that exposure to a novel context evokes glutamate release in the NAcore.

Amperometric measures of NO have been performed previously (Barbosa *et al*. 2008; Ferreira *et al*. 2005). However, to the best of our knowledge we are the first to perform measurements of NO release in behaving animals. While amperometric detection of NO in behaving animals is a new field, citrulline, a coproduct of NO synthesis, can be measured with microdialysis (Saul’skaya *et al*. 2008). Our results are in general agreement with others demonstrating that exposure of rats to a novel context increases citrulline in the NAcore by ∼50% (Saul’skaya & Sudorgina 2014). Moreover, we show here that exposure to a novel context increased NO in the NAcore within 3 minutes, 1 minutes behind what was observed in identical glutamate recordings. Our data support the hypothesis that exposure to a novel context evokes glutamate release, leading to the induction of NO release in the NAcore. Both glutamate and NO are likely involved in the encoding and recognition of novel environmental stimuli, despite the lack of reward availability, yet further experimentation is required to dissect the precise role of each analyte.

### Cue-induced reinstatement and glutamate release in the NAcore

Potentiated glutamate release in the NAcore mediates both cue (Smith *et al*. 2017) and cocaine prime-induced (McFarland *et al*. 2003) reinstatement of cocaine seeking. Glutamate overflow due to enhancement of pre-synaptic glutamate release and decreased glutamate clearance (Baker *et al*. 2002; Xi *et al*. 2002; Baker *et al*. 2003b; Baker *et al*. 2003a; Stefanik & Kalivas 2013) is thought to engage adaptations in MSN structural and synaptic plasticity which drive relapse through NO- and MMP-mediated signaling. This cue-induced relapse signaling cascade then reaches MSNs through integrin receptor activation (Garcia-Keller *et al*. 2019), resulting in transient increases AMPA/NMDA ratios and dendritic spine head diameter of NAcore MSNs. These transient changes in synaptic plasticity are the most profound during the first 15 minutes of reinstatement (Gipson *et al*. 2014; Gipson *et al*. 2013). Measuring glutamate overflow in the NAcore during reinstatement (Smith *et al*. 2017; McFarland *et al*. 2003) has typically been performed using microdialysis (Drew *et al*. 2004). While highly sensitive, microdialysis lacks the temporal resolution afforded by amperometry (Rutherford *et al*. 2007). However, amperometry is well-suited for the near real time detection of evoked glutamate release but is generally considered to be less suitable for the detection of basal glutamate levels. Fortunately, basal levels of NAcore glutamate have been previously established by a body of work using a technique specifically designed for this type of analysis, no-net-flux microdialysis. The general consensus from this body of work places extracellular glutamate concentrations in the NAcore at 1-2 µM (Pati *et al*. 2016; Griffin *et al*. 2015; Moussawi *et al*. 2011). Our amperometric analyses of cue-induced glutamate release in the NAcore yielded an increase of ∼1 μM in cocaine SA rats. When basing a probable basal concentration of glutamate in the NAcore off of the no-net-flux work of 1-2 μM, our data would correspond to a 50-100% increase in extracellular glutamate in the NAcore during relapse. These results fit exceptionally well with previous microdialysis-based estimates of glutamate release in the NAcore during cue-induced reinstatement of cocaine seeking (Smith *et al*. 2017), making our data a sound replication of this original finding.

Interestingly, we found that animals that self-administered saline also show a discernible increase in glutamate release (∼250 nM) during cue-induced reinstatement, despite the fact that saline SA animals showed no clear motivation to seek saline (i.e. lack of significant reinstatement behavior when compared to extinction). While a quantifiable glutamate response was observed in saline SA animals, this elevation in glutamate levels was short-lived. Further, saline SA rats showed a decrease in glutamate below baseline as a function of time during reinstatement. Given the dysfunction in astrocytic-mediated glutamate uptake in the NAcore following extinction from cocaine SA, including retraction of astrocyte processes from synapses (Scofield *et al*. 2016b; Scofield & Kalivas 2014; Scofield *et al*. 2016a; Scofield *et al*. 2015), we posit that the sustained cue-evoked glutamate release in the NAcore in cocaine SA rats is due in part to cocaine-mediated presynaptic potentiation and dysfunctional glutamate transport, which would not be present in saline SA rats. When analyzing the first 15 minutes of reinstatement, we found that glutamate is increased relative to a home-cage baseline in as little as two minutes into cue-induced reinstatement. Interestingly, this is evident in both cocaine and saline SA rats. When comparing glutamate levels during reinstatement, animals reinstating to cocaine cues show a significant elevation of glutamate release beyond what was observed in saline controls 10 minutes into the reinstatement session. This is surprising as this finding indicates that the first 10 minutes of reinstatement and cue exposure is associated with elevated glutamate release in the NAcore regardless of whether the context and cues are associated with cocaine availability.

### Cue-induced reinstatement and NO release in the NAcore

We demonstrate here that cue-induced reinstatement of cocaine seeking elevated NO in the NAcore, with a slightly delayed temporal signature relative to cued reinstatement-evoked glutamate release in the NAcore. Akin to what was observed with glutamate, saline SA rats also displayed a transient increase in NO in the NAcore that decayed as a function of time during reinstatement. NO signaling is implicated in the pathophysiology underlying numerous neuropsychiatric disorders, including psychostimulant addiction (Pulvirenti *et al*. 1996; Hirst & Robson 2011). As an example, nNOS-deficient homozygous mice display a gross impairment in the development and expression of cocaine-induced locomotor sensitization (Itzhak *et al*. 1998a) and a blunted ability to maintain or reinstate cocaine conditioned place preference (CPP) (Balda *et al*. 2006; Itzhak & Anderson 2007). This effect is likely localized to the NAcore given that optogenetic stimulation or inhibition of somatostatin-expressing interneurons in the NAcore, which co-express nNOS and neuropeptide Y (NPY), potentiates or suppresses cocaine CPP, respectively (Ribeiro *et al*. 2018) and nNOS interneuron activation in the NAcore is both necessary and sufficient for cue-induced cocaine seeking (Smith *et al*. 2017). As we have previously shown that somatostatin receptor inhibition in the NAcore is without an effect on cue-induced reinstatement, and that activation of NAcore nNOS interneurons is necessary and sufficient for cue-induced reinstatement (Smith *et al*. 2017), we posit that release of NO from these interneurons is required for relapse.

NO signaling has long been recognized as a vital aspect of plasticity at excitatory synapses, regulating experience- and context-dependent learning (Bon *et al*. 1992). In the striatum, the activation of nNOS interneurons by glutamate-releasing cortical afferents regulates the induction of striatal LTP and LTD (Centonze *et al*. 2003), dysfunction of which has been mechanistically linked to relapse vulnerability (Moussawi *et al*. 2011). Recent monosynaptic tracing experiments have identified the BLA, PL cortex, and ventral subiculum (vSub) as major direct monosynaptic inputs to the nNOS/SOM/NPY class of GABAergic interneurons in the NAcore (Ribeiro *et al*. 2019). Although it is unknown which glutamatergic inputs to nNOS interneurons in the NAcore mediate glutamate-induced NO release, we speculate that the PL cortex is a major contributor given the wealth of data indicating a major role for PL cortical inputs to the NAcore reinstatement of cocaine seeking evoked by multiple stimuli (Scofield *et al*. 2016a). Moreover, glutamatergic cortical inputs are also required for the t-SP in NAcore MSNs linked to the activation of striatal motor programs mediating cue-induced seeking (Gipson *et al*. 2013). Finally, systemic injections of 7-NI decreases striatal NO efflux following electrical stimulation of the frontal cortex of anesthetized rats (Sammut *et al*. 2007), suggesting that corticostriatal glutamate transmission engages nNOS interneurons to drive NO release. Future studies will evaluate the relative contribution of each of these inputs in cue-induced NO release in the NAcore using pathway-specific chemogenetics combined with glutamate and NO amperometry.

Although a clear role for NO-induced activation of MMPs, which are required for t-SP and relapse, via S-nitrosylation has been described (Smith *et al*. 2014; Smith *et al*. 2017), it is also likely that NO release from nNOS interneurons directly engages cellular signaling cascades within MSNs. As an example, S-nitrosylation of N-ethylmaleimide sensitive factor (NSF) (Huang *et al*. 2005), stargazin (Selvakumar *et al*. 2009), as well as GluA1 (Selvakumar *et al*. 2013) also enhances membrane expression of AMPA receptors. Given that cue-induced t-SP is associated with elevated AMPA/NMDA ratios in NAcore MSNs (Gipson *et al*. 2013), it is likely that coordinated activation of both extracellular MMPs as well as AMPA receptor trafficking proteins in NAcore MSNs is required for cue-induced seeking. It should also be noted that more canonical forms of NO signaling, such as increased cGMP through activation of the receptor for NO sGC, may also contribute to the transient alteration of MSN physiological properties driving reinstatement (Lin *et al*. 2010). However, hypotheses regarding additional actions of NO in the NAcore relevant to relapse have yet to be addressed experimentally.

### Dissociation between cued reinstatement-induced and novel context-induced glutamate and NO release in the NAcore

Exposure to a novel context increased glutamate by ∼500 nM in drug-naïve rats, whereas cue-induced reinstatement of cocaine seeking increased glutamate by ∼1 μM. In contrast, novel context-induced NO peaked at ∼20 nM, whereas cue-induced NO peaked at ∼10 nM, in the NAcore, and both increased at the same rate relative to the baseline. In summary, novel context exposure elicits an increase in NAcore glutamate that is of a lower magnitude relative to cued reinstatement-induced glutamate release, yet increased NO magnitude compared to cued reinstatement (Figure S1). Thus, there are likely distinct inputs to the NAcore nNOS neurons responsible for this differential increase in evoked NO levels during exposure to these two distinct contextual stimuli. This also suggests that there is not a simple linear relationship between the extent of glutamate release in the NAcore and the subsequent NO response. Apart from glutamatergic afferents, dopaminergic afferents in the striatum also directly innervate nNOS interneurons (Kerkerian *et al*. 1986; Ribeiro *et al*. 2019). Accordingly, striatal nNOS interneurons express mRNA for dopamine D1 receptors (Le Moine *et al*. 1991). Further, activation of D1 receptors on nNOS interneurons is likely an important regulator of NO release given that systemic injections of D1/5 agonists increases striatal NO efflux in an nNOS-dependent manner (Sammut *et al*. 2006). In addition, glutamatergic inputs that directly synapse on NAcore nNOS neurons, including the BLA (Jones *et al*. 2010) and vSub (Blaha *et al*. 1997), can modulate terminal dopamine release in the NAcore. Further, agonism of D1 receptors has been shown to potentiate NO efflux in the striatum following electrical stimulation of the cortex (Park & West 2009). Thus, it is likely that local glutamate and dopamine interactions in the striatum engage NO efflux to a greater degree than glutamate alone, which may explain the apparent dichotomy between amounts of cue-evoked glutamate and NO relative to the amounts of novel context-evoked glutamate and NO observed herein.

Several lines of evidence indicate that cue-induced reinstatement of cocaine seeking following a short access (2-hour), FR1 schedule of reinforcement SA, extinction, and cue-induced reinstatement paradigm requires elevated glutamate, and not dopamine, transmission in the NAcore (see (Scofield *et al*. 2016a) for review). For example, cue-induced reinstatement is unaffected by blockade of dopamine receptors in the NAcore, yet is prevented by AMPA receptor antagonism (Cornish & Kalivas 2000). In contrast, studies utilizing a second-order schedule of reinforcement have indicated a role for dopamine transmission in the dorsolateral caudate putamen, but not the NAcore, in mediating cocaine seeking by engaging striato-nigro-striatal “spiraling” circuitry (Vanderschuren *et al*. 2005; Belin & Everitt 2008). Importantly, striato-nigral projections are not required for cocaine plus cue-induced reinstatement of cocaine seeking following the SA paradigm used herein (Stefanik *et al*. 2013), suggesting that serial connectivity between the ventral and dorsal striatum is not required for cue-induced reinstatement under an FR1 schedule of reinforcement during SA. Additionally, chemogenetic inhibition of VTA dopamine neurons has no impact on cue-induced reinstatement (Mahler *et al*. 2019). In contrast, exploration of a novel environment elevates dopamine in the NAcore, which is prevented by inactivation of the vSub (Legault & Wise 2001). Not only does the vSub innervate VTA dopamine neuron axon terminals (Blaha *et al*. 1997), but as described above vSub neurons directly synapse on nNOS interneurons in the NAcore (Ribeiro *et al*. 2019; Kraus & Prast 2002). Thus, we posit that the large NO response in the NAcore observed during novel context exposure relative to cue-induced reinstatement may be due to enhanced dopamine transmission in concert with glutamate release, despite the overall decrease in the magnitude of glutamate release observed during novel context exploration. This hypothesis is further supported by the finding that saline SA animals displayed lower evoked NO release during reinstatement compared to both cocaine animals as well as drug-naïve animals exposed to a novel context. This is likely due to the animal’s habituation to the environment and exposure to cues predicting a neutral stimulus, i.e., an i.v. infusion of saline.

## Conclusions

In summary, exposure to a familiar, salient, cocaine-paired context with cues available, leads to a fast onset sustained glutamate and NO response. However, exposure to a familiar but relatively neutral environment, such as a saline-paired context with cues available, produces a transient increase in glutamate and NO, likely due to habituation to the context and lack of salient cues. Together these data demonstrate that the onset of glutamate and NO release in the NAcore is time-dependent and varies given the context and the stimuli associated with that particular context. As expected, cue-induced reinstatement of cocaine seeking increased glutamate and NO release in the NAcore. Surprisingly, exposure of drug-naïve animals to a novel context produced approximately 50% of the glutamate and 200% of the NO response relative to cue-induced reinstatement. Interestingly, despite the differences in the levels of glutamate and NO release in reinstatement and novel context exploration, in both cases we observed a slower onset of NO relative to Glutamate. Given the role of nNOS interneurons in regulating multiple facets of both physiological and pathological processes governing learning and memory (Hirst & Robson 2011), our data contribute to a better understanding of this small population of interneurons and their role in regulating the synaptic plasticity underlying cue-reward associations. Our findings describing the temporal nature of cued reinstatement evoked glutamate release and its relationship to NO release, set the stage for additional investigation of the circuits and cell types required for NO release in the NAcore during cued reinstatement of cocaine seeking.

## Abbreviations

SA: (self-administration)
NO: (nitric oxide)
MSNs: (medium spiny neurons)
nNOS: (neuronal nitric oxide synthase)
NAcore: (nucleus accumbens core)
PL: (prelimbic)
vSub: (ventral subiculum)
VTA: (ventral tegmental area)
GLT-1: (glutamate transporter-1)
BLA: (basolateral amygdala)
DETA: (DETA-NONOate)
CNO: (Clozapine-n-oxide)
FR: (Fixed ratio)
(SEM): standard error of the mean
eGFP: (enhanced green fluorescent protein)
cGMP: (cyclic GMP)
sGC: (soluble guanylyl cyclase)
CPP: (conditioned place preference)
MMPs: (matric metallo-proteinases)
i.v.: (intravenous)
LOD: (limit of detection)
GluOx: (glutamate oxidase)

## Acknowledgements

This work was supported by R00 DA040004 (MDS) and T32 DA007288 (BMS). The authors have no conflicts of interest to disclose.

---

**Figure S1.**
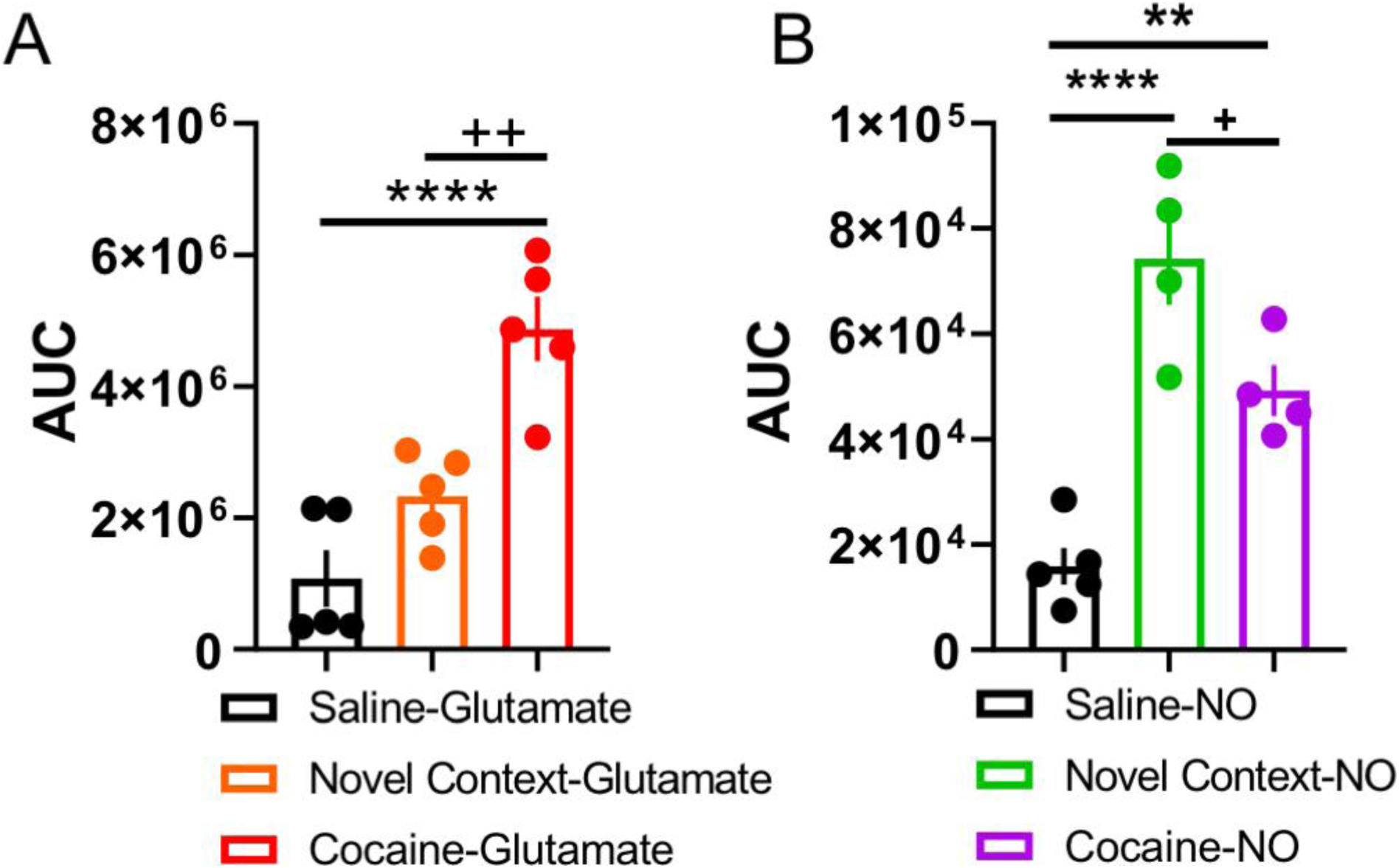
Comparison of the area under the curve (AUC) for the full two-hour session in glutamate and NO animals from experiments 3 and 4. **A**. Cocaine SA animals undergoing glutamate recordings during cue-induced reinstatement in Experiment 4 displayed significantly greater AUC than saline SA animals (Experiment 4) and novel context-exposed rats (Experiment 3). **B**. Novel context-exposed rats (Experiment 3) and cocaine SA rats (Experiment 4) undergoing NO recordings during cue-induced reinstatement displayed a significantly greater AUC than saline SA rats (Experiment 4), yet novel context-exposed rats had a significantly greater AUC than cocaine SA rats. \*\**p*<0.01, \*\*\*\**p*<0.0001 compared to saline SA. +*p*<0.05, ++*p*<0.01 compared to novel context.

## References

Atalla, A. and Kuschinsky, K. (2006) Effects of blockade of glutamate NMDA receptors or of NO synthase on the development or the expression of associative or non-associative sensitization to locomotor activation by morphine. J Neural Transm (Vienna) 113, 1–10.

Babbedge, R. C., Bland-Ward, P. A., Hart, S. L. and Moore, P. K. (1993) Inhibition of rat cerebellar nitric oxide synthase by 7-nitro indazole and related substituted indazoles. Br J Pharmacol 110, 225–228.

Baker, D. A., McFarland, K., Lake, R. W., Shen, H., Tang, X. C., Toda, S. and Kalivas, P. W. (2003a) Neuroadaptations in cystine-glutamate exchange underlie cocaine relapse. Nat Neurosci 6, 743–749.

Baker, D. A., McFarland, K., Lake, R. W., Shen, H., Toda, S. and Kalivas, P. W. (2003b) N-acetyl cysteine-induced blockade of cocaine-induced reinstatement. Ann N Y Acad Sci 1003, 349–351.

Baker, D. A., Shen, H. and Kalivas, P. W. (2002) Cystine/glutamate exchange serves as the source for extracellular glutamate: modifications by repeated cocaine administration. Amino Acids 23, 161–162.

Balda, M. A., Anderson, K. L. and Itzhak, Y. (2006) Adolescent and adult responsiveness to the incentive value of cocaine reward in mice: role of neuronal nitric oxide synthase (nNOS) gene. Neuropharmacology 51, 341–349.

Barbosa, R. M., Lourenco, C. F., Santos, R. M., Pomerleau, F., Huettl, P., Gerhardt, G. A. and Laranjinha, J. (2008) In vivo real-time measurement of nitric oxide in anesthetized rat brain. Methods Enzymol 441, 351–367.

Belin, D. and Everitt, B. J. (2008) Cocaine seeking habits depend upon dopamine-dependent serial connectivity linking the ventral with the dorsal striatum. Neuron 57, 432–441.

Blaha, C. D., Yang, C. R., Floresco, S. B., Barr, A. M. and Phillips, A. G. (1997) Stimulation of the ventral subiculum of the hippocampus evokes glutamate receptor-mediated changes in dopamine efflux in the rat nucleus accumbens. Eur J Neurosci 9, 902–911.

Bon, C., Bohme, G. A., Doble, A., Stutzmann, J. M. and Blanchard, J. C. (1992) A Role for Nitric Oxide in Long-term Potentiation. Eur J Neurosci 4, 420–424.

Bossert, J. M., Marchant, N. J., Calu, D. J. and Shaham, Y. (2013) The reinstatement model of drug relapse: recent neurobiological findings, emerging research topics, and translational research. Psychopharmacology (Berl) 229, 453–476.

Burmeister, J. J. and Gerhardt, G. A. (2001) Self-referencing ceramic-based multisite microelectrodes for the detection and elimination of interferences from the measurement of L-glutamate and other analytes. Anal Chem 73, 1037–1042.

Centonze, D., Gubellini, P., Bernardi, G. and Calabresi, P. (1999) Permissive role of interneurons in corticostriatal synaptic plasticity. Brain Res Brain Res Rev 31, 1–5.

Centonze, D., Gubellini, P., Pisani, A., Bernardi, G. and Calabresi, P. (2003) Dopamine, acetylcholine and nitric oxide systems interact to induce corticostriatal synaptic plasticity. Rev Neurosci 14, 207–216.

Cornish, J. L. and Kalivas, P. W. (2000) Glutamate transmission in the nucleus accumbens mediates relapse in cocaine addiction. J Neurosci 20, RC89.

Drew, K. L., Pehek, E. A., Rasley, B. T., Ma, Y. L. and Green, T. K. (2004) Sampling glutamate and GABA with microdialysis: suggestions on how to get the dialysis membrane closer to the synapse. J Neurosci Methods 140, 127–131.

Epstein, D. H., Preston, K. L., Stewart, J. and Shaham, Y. (2006) Toward a model of drug relapse: an assessment of the validity of the reinstatement procedure. Psychopharmacology (Berl) 189, 1–16.

Ferreira, N. R., Ledo, A., Frade, J. G., Gerhardt, G. A., Laranjinha, J. and Barbosa, R. M. (2005) Electrochemical measurement of endogenously produced nitric oxide in brain slices using Nafion/o-phenylenediamine modified carbon fiber microelectrodes. Analytica Chimica Acta 535, 1–7.

Forstermann, U. and Sessa, W. C. (2012) Nitric oxide synthases: regulation and function. Eur Heart J 33, 829-837, 837a-837d.

Friedemann, M. N., Robinson, S. W. and Gerhardt, G. A. (1996) o-Phenylenediamine-modified carbon fiber electrodes for the detection of nitric oxide. Anal Chem 68, 2621–2628.

Gabbott, P. L., Warner, T. A., Jays, P. R., Salway, P. and Busby, S. J. (2005) Prefrontal cortex in the rat: projections to subcortical autonomic, motor, and limbic centers. J Comp Neurol 492, 145–177.

Garcia-Keller, C., Neuhofer, D., Bobadilla, A. C., Spencer, S., Chioma, V. C., Monforton, C. and Kalivas, P. W. (2019) Extracellular Matrix Signaling Through beta3 Integrin Mediates Cocaine Cue-Induced Transient Synaptic Plasticity and Relapse. Biol Psychiatry 86, 377–387.

Gass, J. T., Sinclair, C. M., Cleva, R. M., Widholm, J. J. and Olive, M. F. (2011) Alcohol-seeking behavior is associated with increased glutamate transmission in basolateral amygdala and nucleus accumbens as measured by glutamate-oxidase-coated biosensors. Addict Biol 16, 215–228.

Giannotti, G., Barry, S. M., Siemsen, B. M., Peters, J. and McGinty, J. F. (2018) Divergent Prelimbic Cortical Pathways Interact with BDNF to Regulate Cocaine-seeking. J Neurosci 38, 8956–8966.

Gipson, C. D., Kupchik, Y. M. and Kalivas, P. W. (2014) Rapid, transient synaptic plasticity in addiction. Neuropharmacology 76 Pt B, 276–286.

Gipson, C. D., Kupchik, Y. M., Shen, H., Reissner, K. J., Thomas, C. A. and Kalivas, P. W. (2013) Relapse induced by cues predicting cocaine depends on rapid, transient synaptic potentiation. Neuron 77, 867–872.

Goren, M. Z., Aricioglu-Kartal, F., Yurdun, T. and Uzbay, I. T. (2001) Investigation of extracellular L-citrulline concentration in the striatum during alcohol withdrawal in rats. Neurochem Res 26, 1327–1333.

Griffin, W. C., Ramachandra, V. S., Knackstedt, L. A. and Becker, H. C. (2015) Repeated cycles of chronic intermittent ethanol exposure increases basal glutamate in the nucleus accumbens of mice without affecting glutamate transport. Front Pharmacol 6, 27.

Gu, Z., Kaul, M., Yan, B., Kridel, S. J., Cui, J., Strongin, A., Smith, J. W., Liddington, R. C. and Lipton, S. A. (2002) S-nitrosylation of matrix metalloproteinases: signaling pathway to neuronal cell death. Science 297, 1186–1190.

Hamel, L., Thangarasa, T., Samadi, O. and Ito, R. (2017) Caudal Nucleus Accumbens Core Is Critical in the Regulation of Cue-Elicited Approach-Avoidance Decisions. eNeuro 4.

Hascup, E. R., Hascup, K. N., Stephens, M., Pomerleau, F., Huettl, P., Gratton, A. and Gerhardt, G. A. (2010) Rapid microelectrode measurements and the origin and regulation of extracellular glutamate in rat prefrontal cortex. J Neurochem 115, 1608–1620.

Hirst, D. G. and Robson, T. (2011) Nitric oxide physiology and pathology. Methods Mol Biol 704, 1–13.

Hu, Y., Mitchell, K. M., Albahadily, F. N., Michaelis, E. K. and Wilson, G. S. (1994) Direct measurement of glutamate release in the brain using a dual enzyme-based electrochemical sensor. Brain Res 659, 117–125.

Huang, Y., Man, H. Y., Sekine-Aizawa, Y. et al. (2005) S-nitrosylation of N-ethylmaleimide sensitive factor mediates surface expression of AMPA receptors. Neuron 46, 533–540.

Isherwood, S. N., Robbins, T. W., Dalley, J. W. and Pekcec, A. (2018) Bidirectional variation in glutamate efflux in the medial prefrontal cortex induced by selective positive and negative allosteric mGluR5 modulators. J Neurochem 145, 111–124.

Itzhak, Y., Ali, S. F., Martin, J. L., Black, M. D. and Huang, P. L. (1998a) Resistance of neuronal nitric oxide synthase-deficient mice to cocaine-induced locomotor sensitization. Psychopharmacology (Berl) 140, 378–386.

Itzhak, Y. and Anderson, K. L. (2007) Memory reconsolidation of cocaine-associated context requires nitric oxide signaling. Synapse 61, 1002–1005.

Itzhak, Y., Martin, J. L., Black, M. D. and Huang, P. L. (1998b) The role of neuronal nitric oxide synthase in cocaine-induced conditioned place preference. Neuroreport 9, 2485–2488.

Jones, J. L., Day, J. J., Aragona, B. J., Wheeler, R. A., Wightman, R. M. and Carelli, R. M. (2010) Basolateral amygdala modulates terminal dopamine release in the nucleus accumbens and conditioned responding. Biol Psychiatry 67, 737–744.

Kalivas, P. W. (2008) Addiction as a pathology in prefrontal cortical regulation of corticostriatal habit circuitry. Neurotox Res 14, 185–189.

Kalivas, P. W. and Volkow, N. D. (2011) New medications for drug addiction hiding in glutamatergic neuroplasticity. Mol Psychiatry 16, 974–986.

Kaneda, T., Sasaki, N., Urakawa, N. and Shimizu, K. (2016) Endothelium-Dependent and -Independent Vasodilator Effects of Dimethyl Sulfoxide in Rat Aorta. Pharmacology 97, 171–176.

Kau, K. S., Madayag, A., Mantsch, J. R., Grier, M. D., Abdulhameed, O. and Baker, D. A. (2008) Blunted cystine-glutamate antiporter function in the nucleus accumbens promotes cocaine-induced drug seeking. Neuroscience 155, 530–537.

Kerkerian, L., Bosler, O., Pelletier, G. and Nieoullon, A. (1986) Striatal neuropeptide Y neurones are under the influence of the nigrostriatal dopaminergic pathway: immunohistochemical evidence. Neurosci Lett 66, 106–112.

Knackstedt, L. A., Melendez, R. I. and Kalivas, P. W. (2010) Ceftriaxone restores glutamate homeostasis and prevents relapse to cocaine seeking. Biol Psychiatry 67, 81–84.

Kraus, M. M. and Prast, H. (2002) Involvement of nitric oxide, cyclic GMP and phosphodiesterase 5 in excitatory amino acid and GABA release in the nucleus accumbens evoked by activation of the hippocampal fimbria. Neuroscience 112, 331–343.

Le Moine, C., Normand, E. and Bloch, B. (1991) Phenotypical characterization of the rat striatal neurons expressing the D1 dopamine receptor gene. Proc Natl Acad Sci U S A 88, 4205–4209.

Legault, M. and Wise, R. A. (2001) Novelty-evoked elevations of nucleus accumbens dopamine: dependence on impulse flow from the ventral subiculum and glutamatergic neurotransmission in the ventral tegmental area. Eur J Neurosci 13, 819–828.

Lin, D. T., Fretier, P., Jiang, C. and Vincent, S. R. (2010) Nitric oxide signaling via cGMP-stimulated phosphodiesterase in striatal neurons. Synapse 64, 460–466.

Mahler, S. V., Brodnik, Z. D., Cox, B. M. et al. (2019) Chemogenetic Manipulations of Ventral Tegmental Area Dopamine Neurons Reveal Multifaceted Roles in Cocaine Abuse. J Neurosci 39, 503–518.

McFarland, K., Lapish, C. C. and Kalivas, P. W. (2003) Prefrontal glutamate release into the core of the nucleus accumbens mediates cocaine-induced reinstatement of drug-seeking behavior. J Neurosci 23, 3531–3537.

Moore, P. K., Babbedge, R. C., Wallace, P., Gaffen, Z. A. and Hart, S. L. (1993) 7-Nitro indazole, an inhibitor of nitric oxide synthase, exhibits antinociceptive activity in the mouse without increasing blood pressure. Br J Pharmacol 108, 296–297.

Moussawi, K., Zhou, W., Shen, H., Reichel, C. M., See, R. E., Carr, D. B. and Kalivas, P. W. (2011) Reversing cocaine-induced synaptic potentiation provides enduring protection from relapse. Proc Natl Acad Sci U S A 108, 385–390.

Park, D. J. and West, A. R. (2009) Regulation of striatal nitric oxide synthesis by local dopamine and glutamate interactions. J Neurochem 111, 1457–1465.

Park, W. K., Bari, A. A., Jey, A. R., Anderson, S. M., Spealman, R. D., Rowlett, J. K. and Pierce, R. C. (2002) Cocaine administered into the medial prefrontal cortex reinstates cocaine-seeking behavior by increasing AMPA receptor-mediated glutamate transmission in the nucleus accumbens. J Neurosci 22, 2916–2925.

Pati, D., Kelly, K., Stennett, B., Frazier, C. J. and Knackstedt, L. A. (2016) Alcohol consumption increases basal extracellular glutamate in the nucleus accumbens core of Sprague-Dawley rats without increasing spontaneous glutamate release. Eur J Neurosci 44, 1896–1905.

Pulvirenti, L., Balducci, C. and Koob, G. F. (1996) Inhibition of nitric oxide synthesis reduces intravenous cocaine self-administration in the rat. Neuropharmacology 35, 1811–1814.

Reissner, K. J., Gipson, C. D., Tran, P. K., Knackstedt, L. A., Scofield, M. D. and Kalivas, P. W. (2015) Glutamate transporter GLT-1 mediates N-acetylcysteine inhibition of cocaine reinstatement. Addict Biol 20, 316–323.

Ribeiro, E. A., Nectow, A. R., Pomeranz, L. E., Ekstrand, M. I., Koo, J. W. and Nestler, E. J. (2019) Viral labeling of neurons synaptically connected to nucleus accumbens somatostatin interneurons. PLoS One 14, e0213476.

Ribeiro, E. A., Salery, M., Scarpa, J. R. et al. (2018) Transcriptional and physiological adaptations in nucleus accumbens somatostatin interneurons that regulate behavioral responses to cocaine. Nat Commun 9, 3149.

Rinaldi, A., Romeo, S., Agustin-Pavon, C., Oliverio, A. and Mele, A. (2010) Distinct patterns of Fos immunoreactivity in striatum and hippocampus induced by different kinds of novelty in mice. Neurobiol Learn Mem 94, 373–381.

Rojo, A. I., McBean, G., Cindric, M., Egea, J., Lopez, M. G., Rada, P., Zarkovic, N. and Cuadrado, A. (2014) Redox control of microglial function: molecular mechanisms and functional significance. Antioxid Redox Signal 21, 1766–1801.

Rutherford, E. C., Pomerleau, F., Huettl, P., Stromberg, I. and Gerhardt, G. A. (2007) Chronic second-by-second measures of L-glutamate in the central nervous system of freely moving rats. J Neurochem 102, 712–722.

Sammut, S., Dec, A., Mitchell, D., Linardakis, J., Ortiguela, M. and West, A. R. (2006) Phasic dopaminergic transmission increases NO efflux in the rat dorsal striatum via a neuronal NOS and a dopamine D(1/5) receptor-dependent mechanism. Neuropsychopharmacology 31, 493–505.

Sammut, S., Park, D. J. and West, A. R. (2007) Frontal cortical afferents facilitate striatal nitric oxide transmission in vivo via a NMDA receptor and neuronal NOS-dependent mechanism. J Neurochem 103, 1145–1156.

Santos, R. M., Lourenco, C. F., Gerhardt, G. A., Cadenas, E., Laranjinha, J. and Barbosa, R. M. (2011) Evidence for a pathway that facilitates nitric oxide diffusion in the brain. Neurochem Int 59, 90–96.

Saul’skaya, N. B., Fofonova, N. V. and Savel’ev, S. A. (2008) Glutamatergic regulation of extracellular citrulline levels in the nucleus accumbens during an emotional conditioned reflex. Neurosci Behav Physiol 38, 487–492.

Saul’skaya, N. B. and Sudorgina, P. V. (2014) Mediolateral Gradient of Nitrergic Activation of the Nucleus Accumbens during Investigative Behavior. Neuroscience and Behavioral Physiology 44, 87–92.

Saulskaya, N. and Marsden, C. A. (1995) Extracellular glutamate in the nucleus accumbens during a conditioned emotional response in the rat. Brain Res 698, 114–120.

Scofield, M. D., Boger, H. A., Smith, R. J., Li, H., Haydon, P. G. and Kalivas, P. W. (2015) Gq-DREADD Selectively Initiates Glial Glutamate Release and Inhibits Cue-induced Cocaine Seeking. Biol Psychiatry 78, 441–451.

Scofield, M. D., Heinsbroek, J. A., Gipson, C. D., Kupchik, Y. M., Spencer, S., Smith, A. C., Roberts-Wolfe, D. and Kalivas, P. W. (2016a) The Nucleus Accumbens: Mechanisms of Addiction across Drug Classes Reflect the Importance of Glutamate Homeostasis. Pharmacol Rev 68, 816–871.

Scofield, M. D. and Kalivas, P. W. (2014) Astrocytic dysfunction and addiction: consequences of impaired glutamate homeostasis. Neuroscientist 20, 610–622.

Scofield, M. D., Li, H., Siemsen, B. M., Healey, K. L., Tran, P. K., Woronoff, N., Boger, H. A., Kalivas, P. W. and Reissner, K. J. (2016b) Cocaine Self-Administration and Extinction Leads to Reduced Glial Fibrillary Acidic Protein Expression and Morphometric Features of Astrocytes in the Nucleus Accumbens Core. Biol Psychiatry 80, 207–215.

Selvakumar, B., Huganir, R. L. and Snyder, S. H. (2009) S-nitrosylation of stargazin regulates surface expression of AMPA-glutamate neurotransmitter receptors. Proc Natl Acad Sci U S A 106, 16440–16445.

Selvakumar, B., Jenkins, M. A., Hussain, N. K., Huganir, R. L., Traynelis, S. F. and Snyder, S. H. (2013) S-nitrosylation of AMPA receptor GluA1 regulates phosphorylation, single-channel conductance, and endocytosis. Proc Natl Acad Sci U S A 110, 1077–1082.

Siemsen, B. M., Giannotti, G., McFaddin, J. A., Scofield, M. D. and McGinty, J. F. (2018a) Biphasic effect of abstinence duration following cocaine self-administration on spine morphology and plasticity-related proteins in prelimbic cortical neurons projecting to the nucleus accumbens core. Brain Struct Funct.

Siemsen, B. M., Lombroso, P. J. and McGinty, J. F. (2018b) Intra-prelimbic cortical inhibition of striatal-enriched tyrosine phosphatase suppresses cocaine seeking in rats. Addict Biol 23, 219–229.

Siemsen, B. M., Reichel, C. M., Leong, K. C. et al. (2019) Effects of Methamphetamine Self-Administration and Extinction on Astrocyte Structure and Function in the Nucleus Accumbens Core. Neuroscience 406, 528–541.

Sinha, R. (2011) New findings on biological factors predicting addiction relapse vulnerability. Curr Psychiatry Rep 13, 398–405.

Smith, A. C., Kupchik, Y. M., Scofield, M. D., Gipson, C. D., Wiggins, A., Thomas, C. A. and Kalivas, P. W. (2014) Synaptic plasticity mediating cocaine relapse requires matrix metalloproteinases. Nat Neurosci 17, 1655–1657.

Smith, A. C. W., Scofield, M. D., Heinsbroek, J. A. et al. (2017) Accumbens nNOS Interneurons Regulate Cocaine Relapse. J Neurosci 37, 742–756.

Stefanik, M. T. and Kalivas, P. W. (2013) Optogenetic dissection of basolateral amygdala projections during cue-induced reinstatement of cocaine seeking. Front Behav Neurosci 7, 213.

Stefanik, M. T., Kupchik, Y. M., Brown, R. M. and Kalivas, P. W. (2013) Optogenetic evidence that pallidal projections, not nigral projections, from the nucleus accumbens core are necessary for reinstating cocaine seeking. J Neurosci 33, 13654–13662.

Stefanik, M. T., Kupchik, Y. M. and Kalivas, P. W. (2016) Optogenetic inhibition of cortical afferents in the nucleus accumbens simultaneously prevents cue-induced transient synaptic potentiation and cocaine-seeking behavior. Brain Struct Funct 221, 1681–1689.

Vanderschuren, L. J., Di Ciano, P. and Everitt, B. J. (2005) Involvement of the dorsal striatum in cue-controlled cocaine seeking. J Neurosci 25, 8665–8670.

Wakabayashi, K. T. and Kiyatkin, E. A. (2012) Rapid changes in extracellular glutamate induced by natural arousing stimuli and intravenous cocaine in the nucleus accumbens shell and core. J Neurophysiol 108, 285–299.

Xi, Z. X., Baker, D. A., Shen, H., Carson, D. S. and Kalivas, P. W. (2002) Group II metabotropic glutamate receptors modulate extracellular glutamate in the nucleus accumbens. J Pharmacol Exp Ther 300, 162–171.

Zhang, R., Zhang, L., Zhang, Z., Wang, Y., Lu, M., Lapointe, M. and Chopp, M. (2001) A nitric oxide donor induces neurogenesis and reduces functional deficits after stroke in rats. Ann Neurol 50, 602–611.

